# Genetic basis of plasticity for forage quality traits in response to water deficit in a diverse germplasm panel of alfalfa

**DOI:** 10.1101/554402

**Authors:** Long-Xi Yu, Bill Boge, Jinguo Hu, Steven Fransen, Steven Norberg

**Affiliations:** United States Department of Agriculture-Agricultural Research Service, Plant and Germplasm Introduction and Testing Research; Irrigated Agriculture Extension and Research Center, Washington State University, 24106 N Bunn Road, Prosser, Washington; Washington State University Franklin County Extension Office, 404 West Clark Street, Pasco, Washington, USA

**Author notes:** **Corresponding author**: Long-Xi Yu, United States Department of Agriculture-Agricultural Research Service, 24106 N Bunn Road, Prosser, WA 99350, USA.

**Keywords:** Forage quality, drought, GWA, genetic loci, phenotypic plasticity, alfalfa.

## Abstract

Plant phenotypic plasticity is the ability of plants to express different phenotypes in response to environmental variations. Genetic bases by which phenotypic plasticity affects plant adaptation to environmental change remain largely unknown. In the present study, we characterized 26 forage quality traits in a panel of alfalfa 198 accessions in a field trial under water deficit gradient. The regression analysis revealed that the values of fiber-related traits decreased, while those among energy-related traits increased, as water deficit increased. Genetic loci for forage quality traits were investigated by Genome-wide association studies (GWAS) under different levels of water deficit. Genetic loci associated with forage quality traits were identified and compared. Similar regions were found between energy-related traits when grand means were used for GWAS. Most of the associated markers were identified under water deficit, suggesting genetic mechanisms for forage quality traits were differ between well-watered and water stressed plants. Although GWAS on forage quality have been reported, we are the first to address the genetic factors for forage quality traits under water deficit in autotetraploid alfalfa. The information gained from the present study will be useful for the genetic improvement of alfalfa with enhanced drought/salt tolerance while maintaining forage quality.

## Introduction

Alfalfa (*Medicago sativa* L.), “Queen of the Forages”, is the most productive and highest quality forage crop. Alfalfa quality reflects its potential to produce a desirable animal response and is directly characterized by laboratory measurements of chemical components including protein, fiber and lignin contents, total digestible nutrients, and other calculated parameters such as: relative feed value (RFV), dry matter intake (DMI) potential and relative forage quality (RFQ). Alfalfa quality is important due to its ability to improve animal performance. Fiber contents such as acid detergent fiber (ADF) and neutral detergent fiber (NDF) are important factors affecting rumen function and digestibility of forage. Alfalfa forage contains 35-55% NDF, which contributes ~20-30% of the digestible energy value of alfalfa, the rest coming from non-fiber components (Van Soest, 1982). Relative feed value is a tool that indexes alfalfa quality based on its acid detergent fiber (ADF) and NDF content. The RFV index estimates digestible dry matter (DDM) of the alfalfa from ADF, and calculates the dry matter intake (DMI) potential from NDF. However, RFV has a significant shortcoming because it does not take into account how variations in NDF digestibility affect the energy content or intake potential of alfalfa. Even when harvested at an immature stage, the digestibility of alfalfa fiber can be very different (Goeser and Combs, 2009). In 2004, scientists at the University of Wisconsin designed another index, relative forage quality (RFQ) for estimating forage quality. The RFQ uses fiber digestibility and the total digestible nutrients of the forage to estimate intake (Undersander and Moore, 2004). The RFQ index is an improvement over the RFV index as it better reflects the performance on animal fed. The RFQ emphasizes fiber digestibility while RFV uses digestible dry matter intake. The RFV continues to be widely used as an index to assess quality, compare forage varieties, and price forages. However, differences in the digestibility of the fiber fraction can result in a difference in animal performance when forages with a similar RFV index are fed. The RFQ index has been developed to overcome this difference and it takes into account the differences in digestibility of the fiber fraction and can be used to more accurately predict animal performance.

Alfalfa forage quality is affected by environmental factors. Irrigation and an arid climate affect nutritive value of alfalfa hay. Environmental factors such as drought and high salinity are frequently occurred in the arid and semi-arid regions and affect plant growth. Plants have developed several mechanisms to cope with these challenges: including adapting themselves to survive in the adverse conditions (stress-tolerance), and/or changing growth habits to avoid stress conditions (stress-avoidance). Stress-tolerant plants have evolved certain adaptive mechanisms such as phenotypic plasticity to achieve different degrees of tolerance. The extent of phenotypic plasticity often involves in genetic basis subjected to selective pressure, and can evolve a mechanism that facilitates adaptation to environmental changes (Schlichting, 1986). It is unclear whether plant phenotypic plasticity is controlled by specific genes or a result of epistatic interaction during the selection of individual traits. However, genetic diversity and heterozygosity enhance adaptability to variable environments. It is important to identify plant functional traits by which plasticity may play a causal role in plant response to global climate change (Gratani, 2004) or year to year weather changes.

In the present study, we evaluated 26 forage quality traits in a panel of 198 alfalfa accessions collected from northern America and dryland regions of other countries. The plants were evaluated in the field under three irrigation regimes: well-watered, mild and severe water deficits. To investigate the genetic base of the forage quality, we applied an integrated framework that merges a QTL mapping approach called “genome-wide association studies (GWAS)” with high-throughput genome sequencing methodologies called “genotyping by sequencing (GBS)” to investigate genomic architecture of phenotypic plasticity of alfalfa quality traits under a gradient of water deficits. The ultimate goal is to gain a better understanding on genetic bases by which water deficit affects phenotypic plasticity of forage quality and to identify genetic regions associated with forage quality traits in alfalfa under a deficit irrigation gradient. The identified DNA markers that are significantly associated with the traits will be used for marker-assisted selection and would facilitate breeding for high quality alfalfa varieties based on genetic potential and reduce the confounding of environmental conditions with traditional breeding methods.

## Materials and Methods

### Plant Materials

A panel of germplasm composed of 198 alfalfa accessions selected from the USDA-ARS National Plant Germplasm System (NPGS) alfalfa collection was used in this study. Majority of the panel was collected from Northwest regions of US and Canada including Manitoba, Idaho, Montana, Nebraska, Washington, North and South Dakota. British Columbia, and Saskatchewan and the remaining was collected from dry regions of different countries including Afghanistan, Algeria, Bulgaria, China, Germany, India, Lebanon, Oman, Russia, Spain, Turkey, and Yemen.

### Field experiments

Alfalfa accessions were planted on the Roza Farm (GPS point is 46.289391, −119.725428) at the Irrigated Agriculture Research and Extension Center, Washington State University, Prosser WA, in 2016. Field experiments used a randomized complete block design. Each block contains 9 plants with 15 inch between rows and 10 inch between plants. They were irrigated regularly until the plant size was uniform (day 35-40). Subsequently, the control plants were watered regularly while water deficits were applied to the stressed plants by withholding water during the dry season (June-September). Periodic irrigation was applied as necessary for the mild stress. The exact watering interval was determined during the trial through visual evaluation of wilting of the plants. No water was applied until harvesting for the severe drought stressed plants.

### Forage quality measurements

A subset of samples from the first cuttings was used for quality analyses. Plant samples were dried in oven at 60°C. They were then ground in Wiley Mill (Thomas Scientific, US) prior to the final grinding in Cyclotec 1093 sampling mill (Foss, Hillerød, Denmark) through a 1 mm screen. Sample powders were loaded and measured by Near Infrared Reflectance Spectroscopy (NIRS). Spectra were collected by a scanning monochromator (FOSSNIR Systems 6500, Silver Spring, MD, USA) in the spectral range from 400 to 2500 nm. A NIRS Consortium equation 1 3AH50.2-eqa was used to predict quality factors.

### Statistical analysis

Phenotypic data were subjected to an analysis of variance (ANOVA), testing the variation among accessions for each treatment conditions. To estimate phenotypic plasticity, a plasticity index (PI) was calculated according to Valladares et al., (2000) as follow:

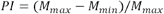

Where *M_max_* is the highest value of the treatment average and *M_min_* is the lowest value of treatment average for a specific trait in the population.

### Genotyping by sequencing

Genotyping-by-sequencing was carried out as described Elshire et al. 2011). Briefly, DNA extracted from original plants of the association panel was digested by methylation sensitive restriction enzyme, EcoT221. GBS libraries were then prepared with barcode adapters. Libraries were sequenced in the Illumina Hi-Seq2000 instrument using 100-base single-end sequencing with 2 lanes at Cornell University Sequencing facility (Ithaca, NY).

After quality check using FastQC (http://www.bioinformatics.babraham.ac.uk/projects/fastqc/), the sequence reads were proceeded using the FreeBayes pipeline adapted for tetraploid alfalfa for genotype calling as described previously (Yu et al. 2017).

### Genome-wide association analysis

The filtered marker data were loaded to TASSEL (Bradbury et al. 2007) for GWAS. A mixed linear model was used for analyzing marker-trait association. The Kinship (K) and Q matrices were used for controlling population structure during association analysis. Marker’s p-value was used to determine the significance of marker-trait association. A false discovery rate (FDR) of 0.05 was used as a threshold for significant association (Benjamini and Hochberg, 1995).

## Results

### Phenotypic variations of forage quality traits

The analysis of variance for 26 quality traits was carried out among the panel of germplasm and the result is presented in Table 1. The sum of squares varied from 0.29 in NEL to 130770.31 in RFV. The differences of most of the traits are statistically significant at the probability <0.0001 level. The estimated parameters varied from 0.41 (NEG) to 164.71 (RFV). The differences for the parameter estimates are statistically significant at the probability <0.0001 level for all the traits. Regressions of phenotypic evaluations showed significant effects of water deficit on forage quality (Fig. 1). The values of fiber-related traits, including ADF, aNDF, dNDF30, dNDF48 decreased as water deficit applied (Fig. 1, A1-4). Water deficit also decreased the contents of fat, RUP, IVDDM30 and NDFD48 (Fig. 1 B1-4), and slightly decreased TDNL, protein, IVDDM48 and lignin (Fig. 1, C1-4). Whereas, the values for energy-related traits include DM, ME, NEL, TDN, NEM, NEG, DRYMI, RFV, ENE, DDM, NFC and DMI1 increased by water deficit (Fig. 1 D1to F4). Water deficit slightly increased RFQ and ash (Fig. 1, G1-2).

**Fig. 1.**
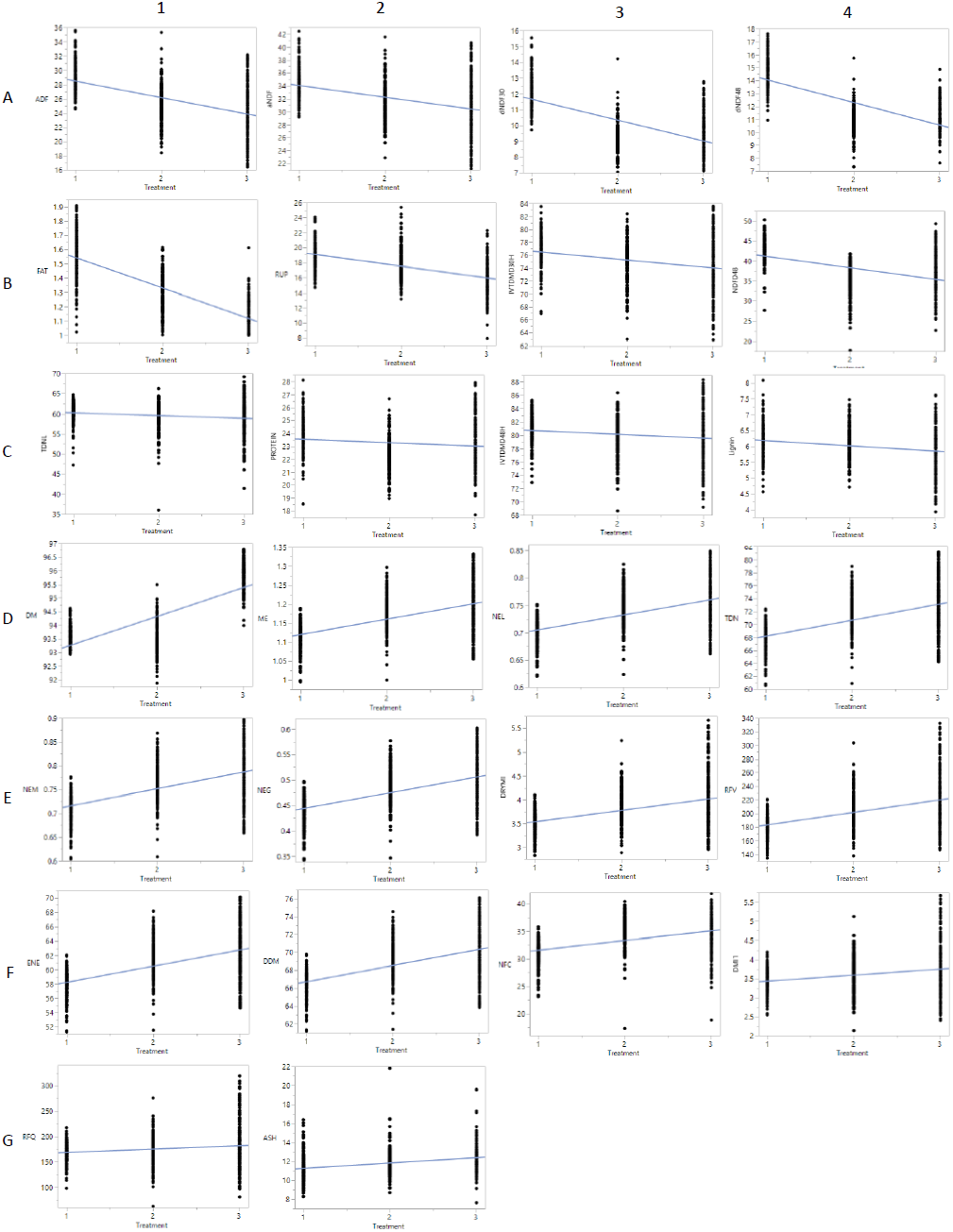
Bivariable plots for forage quality traits under well-watered (1), mild (2) and severe (3) water deficits (X-axis). Phenotypic values of each trait are presented by Y-axis.

**Table 1.** Analysis of variances of forage quality traits in the panel of 198 alfalfa accessions

| Analysis of Variance |  |  |  | Parameter Estimates |  |  |  |
| --- | --- | --- | --- | --- | --- | --- | --- |
| Variable | Sum of Squares | F Ratio | Prob > F | Estimate | Std Error | t Ratio | Prob>\|t\| |
| DM | 437.53 | 711.77 | <.0001 | 92.19 | 0.09 | 1076.20 | <.0001 |
| CP | 26.72 | 9.45 | 0.0022 | 23.81 | 0.18 | 129.60 | <.0001 |
| ADF | 2070.37 | 231.21 | <.0001 | 30.82 | 0.33 | 94.25 | <.0001 |
| aNDF | 1276.96 | 104.09 | <.0001 | 35.90 | 0.38 | 93.80 | <.0001 |
| IVTDMD48 | 125.19 | 13.11 | 0.0003 | 81.29 | 0.34 | 240.76 | <.0001 |
| dNDF48 | 1176.80 | 548.38 | <.0001 | 15.80 | 0.16 | 98.71 | <.0001 |
| IVTDMD30 | 577.79 | 45.29 | <.0001 | 77.70 | 0.39 | 199.10 | <.0001 |
| dNDF30 | 681.32 | 386.22 | <.0001 | 12.99 | 0.15 | 89.54 | <.0001 |
| ASH | 121.61 | 58.16 | <.0001 | 10.69 | 0.16 | 67.68 | <.0001 |
| FAT | 17.27 | 920.86 | <.0001 | 1.75 | 0.01 | 117.13 | <.0001 |
| Lignin | 10.91 | 29.56 | <.0001 | 6.36 | 0.07 | 95.74 | <.0001 |
| RUP | 911.85 | 220.29 | <.0001 | 20.61 | 0.22 | 92.70 | <.0001 |
| NEL | 0.29 | 231.21 | <.0001 | 0.68 | 0.00 | 174.08 | <.0001 |
| TDN | 2364.05 | 231.21 | <.0001 | 65.72 | 0.35 | 188.10 | <.0001 |
| ENE | 2000.33 | 231.21 | <.0001 | 55.94 | 0.32 | 174.08 | <.0001 |
| ME | 0.64 | 231.21 | <.0001 | 1.08 | 0.01 | 188.10 | <.0001 |
| NEM | 0.48 | 230.93 | <.0001 | 0.68 | 0.00 | 136.66 | <.0001 |
| NEG | 0.37 | 230.73 | <.0001 | 0.41 | 0.00 | 93.76 | <.0001 |
| DDM | 1256.39 | 231.21 | <.0001 | 64.90 | 0.25 | 254.80 | <.0001 |
| DRYMI | 22.16 | 114.24 | <.0001 | 3.30 | 0.05 | 68.52 | <.0001 |
| NDFD48 | 3279.66 | 153.65 | <.0001 | 44.12 | 0.50 | 87.40 | <.0001 |
| RFV | 130770.31 | 138.82 | <.0001 | 164.71 | 3.35 | 49.12 | <.0001 |
| NFC | 1158.15 | 121.12 | <.0001 | 29.84 | 0.34 | 88.33 | <.0001 |
| TDNL | 187.28 | 13.66 | 0.0002 | 60.95 | 0.40 | 150.66 | <.0001 |
| DMI1 | 9.74 | 36.29 | <.0001 | 3.27 | 0.06 | 57.82 | <.0001 |
| RFQ | 17203.35 | 14.21 | 0.0002 | 161.78 | 3.80 | 42.56 | <.0001 |

Correlation coefficient was also investigated among the quality traits under well-watered and water deficit conditions. Fig. 2 shows the heat maps of correlations among 26 quality traits under well-watered (Fig. 2 A), mild (Fig. 2 B) and severe (Fig. 2 C) water deficits. Overall, negative correlations were found between DM, ADF, aNDF, ash, lignin, RUP and energy-related traits. Whereas crude protein, IVTDMD30/48 were positively correlated with energy traits. Higher correlations were found among energy traits under all water conditions. The correlations between NDFD48, NFC, TDNL and energy traits increased when water deficits applied.

**Fig. 2.**
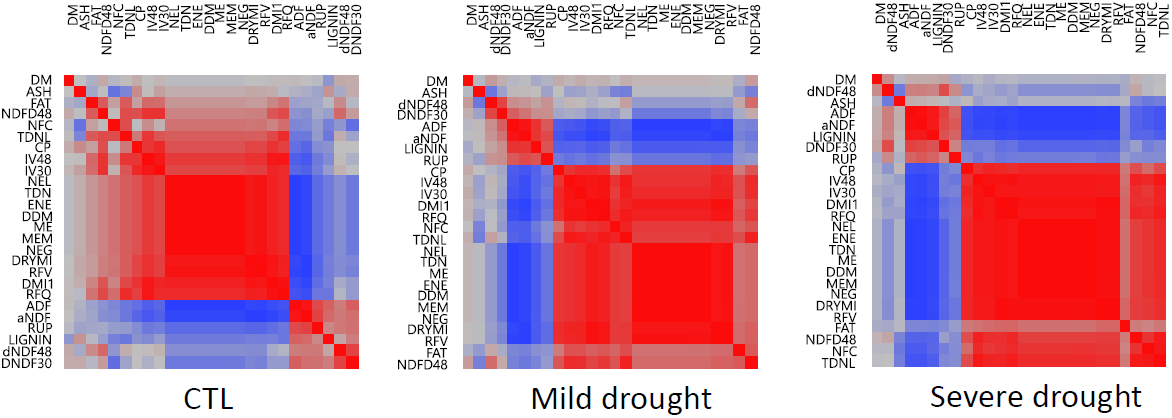
Heat plots of correlation coefficient among quality traits under well-watered control (CTL), mild water deficit (Mild) and severe water deficit (Severe).

### Phenotypic plasticities of quality traits

Phenotypic plasticity was estimated by calculating the PI for each trait in the given accession under well-watered and water deficit conditions as described in the section of Materials and Methods. Overall, higher PIs were found in water deficit conditions compared to well-watered control except lignin, fat and protein (Fig. 3, Table 2). Within the water stress treatments, higher PIs were found in the mild stress than those in the severe stress for most of the quality traits. Among them, the highest PIs were found in RFQ with 0.55, 0.77 and 0.75 for the control, mild and severe drought conditions, respectively (Table 2, Top). The lowest PIs appeared in DM with 0.02, 0.04 and 0.03 for control, mild and severe drought, respectively (Table 2, bottom). The rest of the traits showed higher PIs in severe drought compared to the mild treatment. The PI values were very similar between DRYMI and aNDF, ENE and NEL, and TDN and ME.

**Fig. 3.**
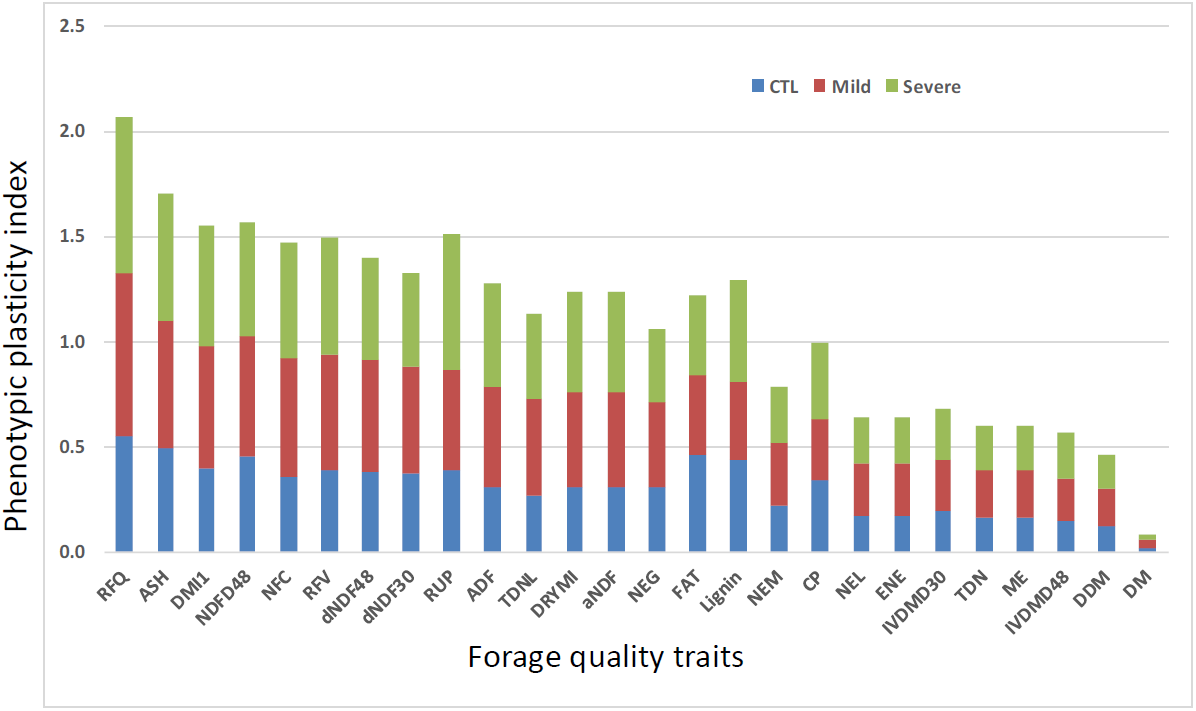
Phenotypic plasticities of 26 forage quality traits in response to mild and severe water deficits and well-watered control in alfalfa.

**Table 2.**
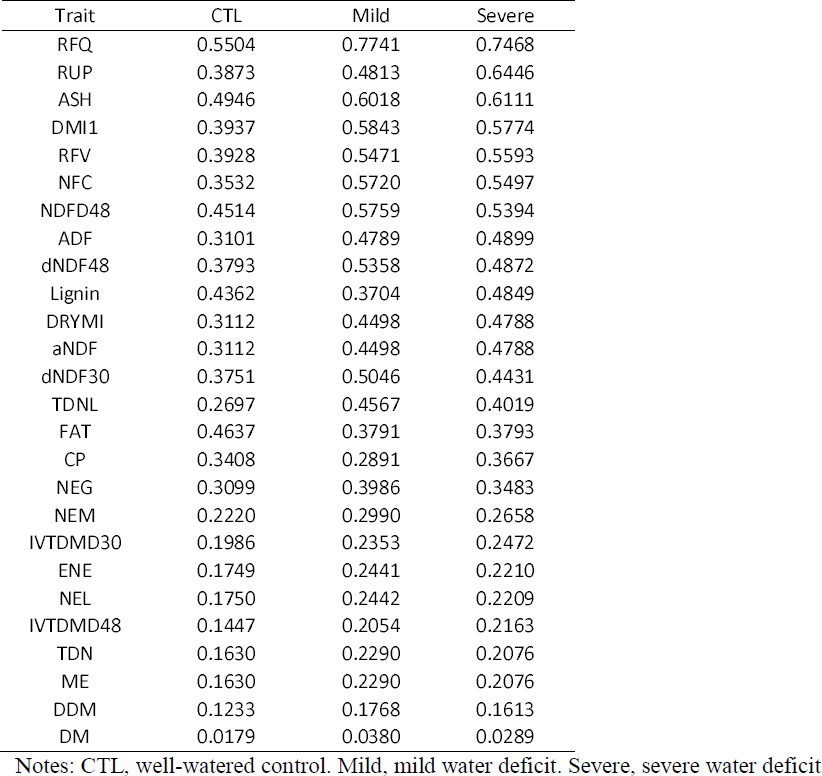
Phenotypic plasticity index for forage quality traits in alfalfa under a water deficit gradient

| Trait | CTL | Mild | Severe |
| --- | --- | --- | --- |
| RFQ | 0.5504 | 0.7741 | 0.7468 |
| RUP | 0.3873 | 0.4813 | 0.6446 |
| ASH | 0.4946 | 0.6018 | 0.6111 |
| DMI1 | 0.3937 | 0.5843 | 0.5774 |
| RFV | 0.3928 | 0.5471 | 0.5593 |
| NFC | 0.3532 | 0.5720 | 0.5497 |
| NDFD48 | 0.4514 | 0.5759 | 0.5394 |
| ADF | 0.3101 | 0.4789 | 0.4899 |
| dNDF48 | 0.3793 | 0.5358 | 0.4872 |
| Lignin | 0.4362 | 0.3704 | 0.4849 |
| DRYMI | 0.3112 | 0.4498 | 0.4788 |
| aNDF | 0.3112 | 0.4498 | 0.4788 |
| dNDF30 | 0.3751 | 0.5046 | 0.4431 |
| TDNL | 0.2697 | 0.4567 | 0.4019 |
| FAT | 0.4637 | 0.3791 | 0.3793 |
| CP | 0.3408 | 0.2891 | 0.3667 |
| NEG | 0.3099 | 0.3986 | 0.3483 |
| NEM | 0.2220 | 0.2990 | 0.2658 |
| IVTDMD30 | 0.1986 | 0.2353 | 0.2472 |
| ENE | 0.1749 | 0.2441 | 0.2210 |
| NEL | 0.1750 | 0.2442 | 0.2209 |
| IVTDMD48 | 0.1447 | 0.2054 | 0.2163 |
| TDN | 0.1630 | 0.2290 | 0.2076 |
| ME | 0.1630 | 0.2290 | 0.2076 |
| DDM | 0.1233 | 0.1768 | 0.1613 |
| DM | 0.0179 | 0.0380 | 0.0289 |

Notes: CTL, well-watered control. Mild, mild water deficit. Severe, severe water deficit

### Cluster analysis of germplasm using forage quality traits

The mean values of 26 forage quality traits were used for cluster analysis. Two large clusters and 14 subclusters were classified as showing in Fig. 4. The first large cluster contained 8 subclusters. Most of germplasm in this cluster were collected from cultivars from US and Canada and their quality traits such as crude protein and RFV were relatively higher, so we named it as the higher forage quality cluster (Fig. 4, top cluster). Two checks, Rambler and Saranac, susceptible to salt/drought are in this cluster (Fig. 4, subclusters 5 and 8). The bottom cluster was furtherly classified into 6 subclusters containing germplasm collected worldwide, including old cultivars and landraces with relatively lower forage quality (Fig. 4, bottom cluster). Three salt/drought resistance checks, Malone, Mesa Sirsa and Wilson are in this cluster (Fig. 4 subcluster 9). There was a trend that alfalfa germplasm with resistance to salt/drought had lower forage quality, while higher quality was found in the susceptible alfalfa germplasm.

**Fig. 4.**
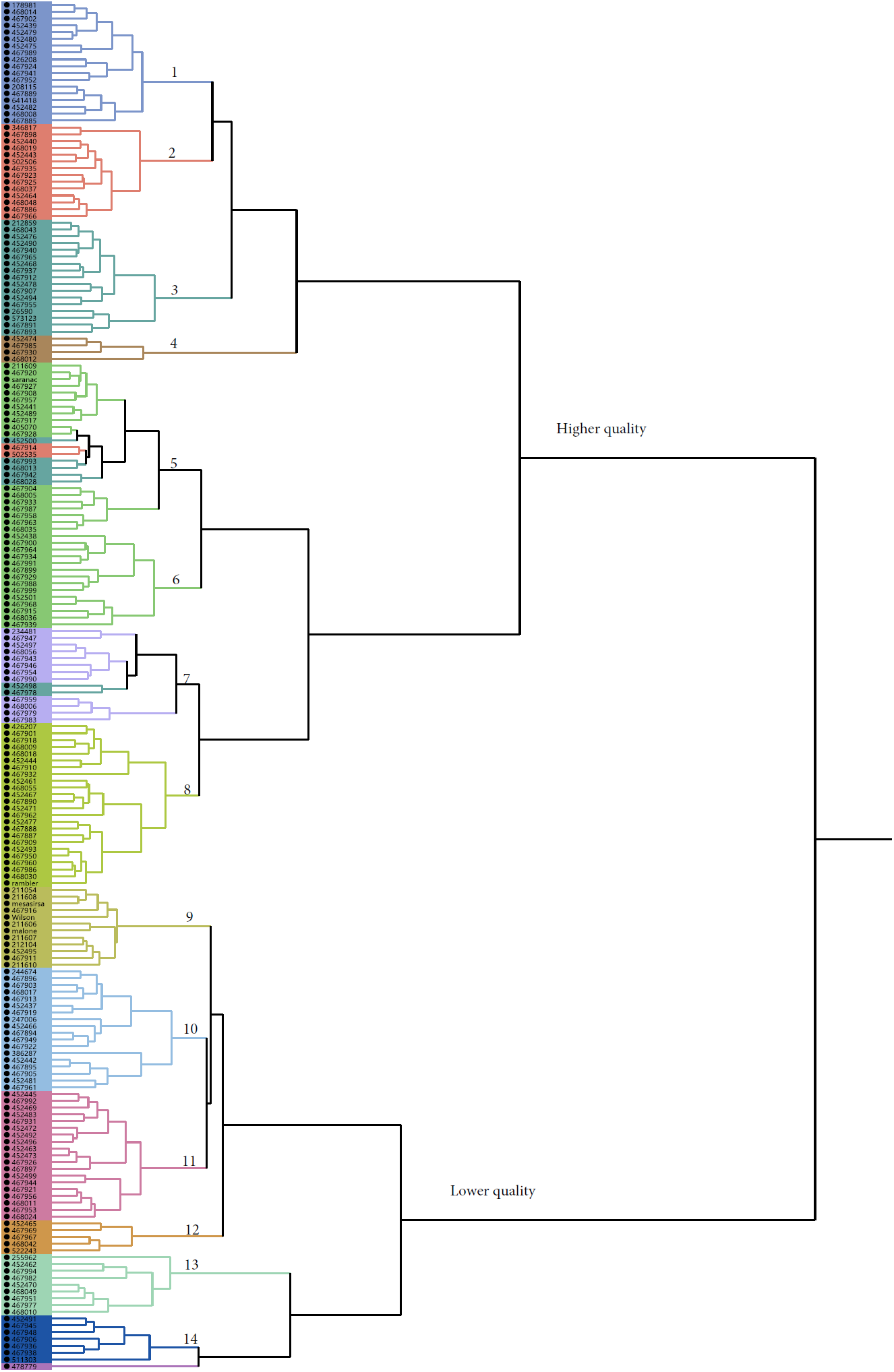
A hierarchical cluster obtained using farthest neighbor method with phenotypic values of all quality traits evaluated in the present study. Accessions (PIs) were clustered into 2 clusters (High and low quality clusters) and 14 subclusters. The high quality cluster contains 8 subclusters with relatively higher quality. The low quality cluster contains 6 subclusters with relatively lower quality. Two checks, “Saranac” and “Rambler”, susceptible to salt/drought stress were clustered into the high quality cluster (Subclusters 5 and 8, respectively), and three drought/salt resistance checks, “Mesa Sirsa”, “Wilson” and “Malone” were clustered into the low quality cluster (Subcluster 9).

### Genome-wide association for forage quality

The combinations of the filtered GBS markers and phenotypic data of 26 quality traits were analyzed by GWAS using a mixed linear model. The results of marker-trait association for well-watered control (A), mild (B) and severe (C) water deficits were illustrated in quantile-quantile plots (QQ) (Fig. 5). The QQ plots shows the expected distributions of association test statistics (X-axis) across thousands of SNPs compared to the observed values (Y-axis). Any deviation from the X=Y line implies a consistent difference between expected and observed across the whole genome. In the present study, as showing in Fig. 5, solid lines represent stress treatments match with the expected line until they sharply curve at the end, representing a small number of true associations among majority of unassociated SNPs. The significances of marker-trait association were presented in negative log P-values. Overall, lower significance was obtained in the control (Fig. 5 A) while higher significances were obtained in the stress (Fig. 5 B and C) with highest in the mild stress Fig. 5 B). Significances were only found in TDNL, NDFD48 and ME under well-watered condition (Fig. 5 A), whereas most significances were found in majority of the traits under the mild stress (Fig. 5 B) with highest in dNDF30 and 48. The highest association was found in ash in the severe stress (Fig. 5 C), however, the level of significances was lower by the severe stress compared to that of the mild stress (Fig. 5 B). More detailed profile of marker-trait association is presented in Manhattan plots (Fig. 6). Of 26 traits analyzed, most significant marker-trait associations were found in ash, NDFD48, dNDF30, dNDF48, NFC and TDNL under mild water stress (Fig. 6 B, E, H, K, N and Q). Similar association profiles were found in the severe stress, but the marker’s significances were lower for the same traits (Fig. 6 C, F, I, L, O and R) compared to the mild stress. Whereas no or less significant markers were shown under control and severe stress conditions (Fig. 6 remaining panels). Among those, the highest significant markers were identified in ash and they were located on chromosomes 2, 6, 7 and unknown chromosome (U) (Fig. 6 B). Similarly, Significant markers associate with NDFD48 were also identified on same chromosomes (Fig. 6 E) under mild stress but not in control and severe stress (Fig. 6 D and F, respectively). Significant markers were also found in dNDF30 and dNDF48 and they were located on chromosomes 1, 2, 3 and 8 (Fig. 6 H and K, respectively). Mild stress also triggered marker-trait associations in NFC and TDNL (Fig. 6 N and Q, respectively). Significant markers associated with NFC were located on chromosomes 2, 6 and 7 (Fig. 6 N). while markers associated with TDNL were located on chromosomes 2 and 6 (Fig. 6 Q).

**Fig. 5.**
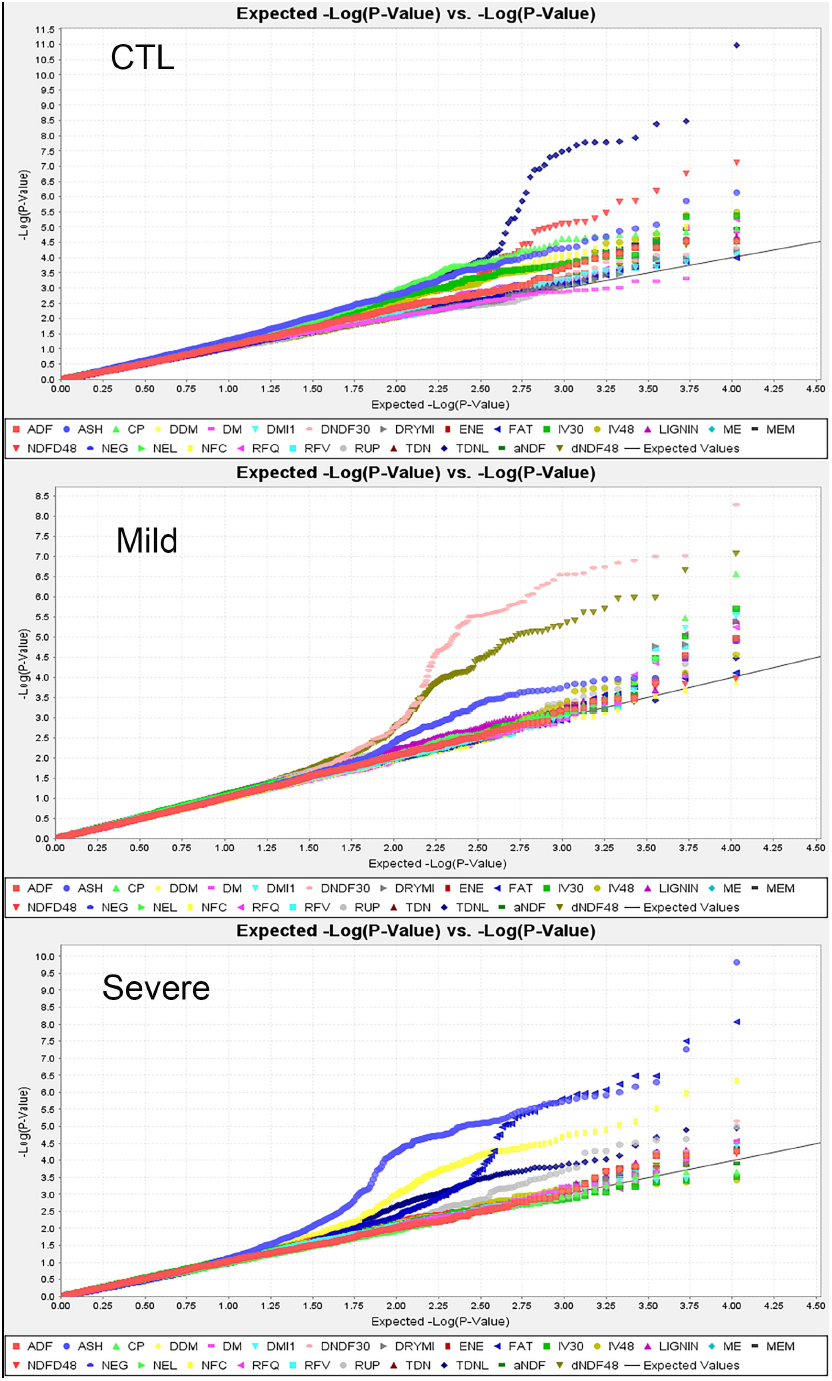
Quantile-quantile plots of marker-trait association from GWAS for forage quality traits under well-watered (A), mild (B) and severe (C) water deficits in the alfalfa association panel. The expected (solid lines) against observed (dot lines) −log_10_ p-values are presented on X-axis and Y-axis, respectively. Each color curve represents a quality trait as showing at the bottom of the figures.

**Fig. 6.**
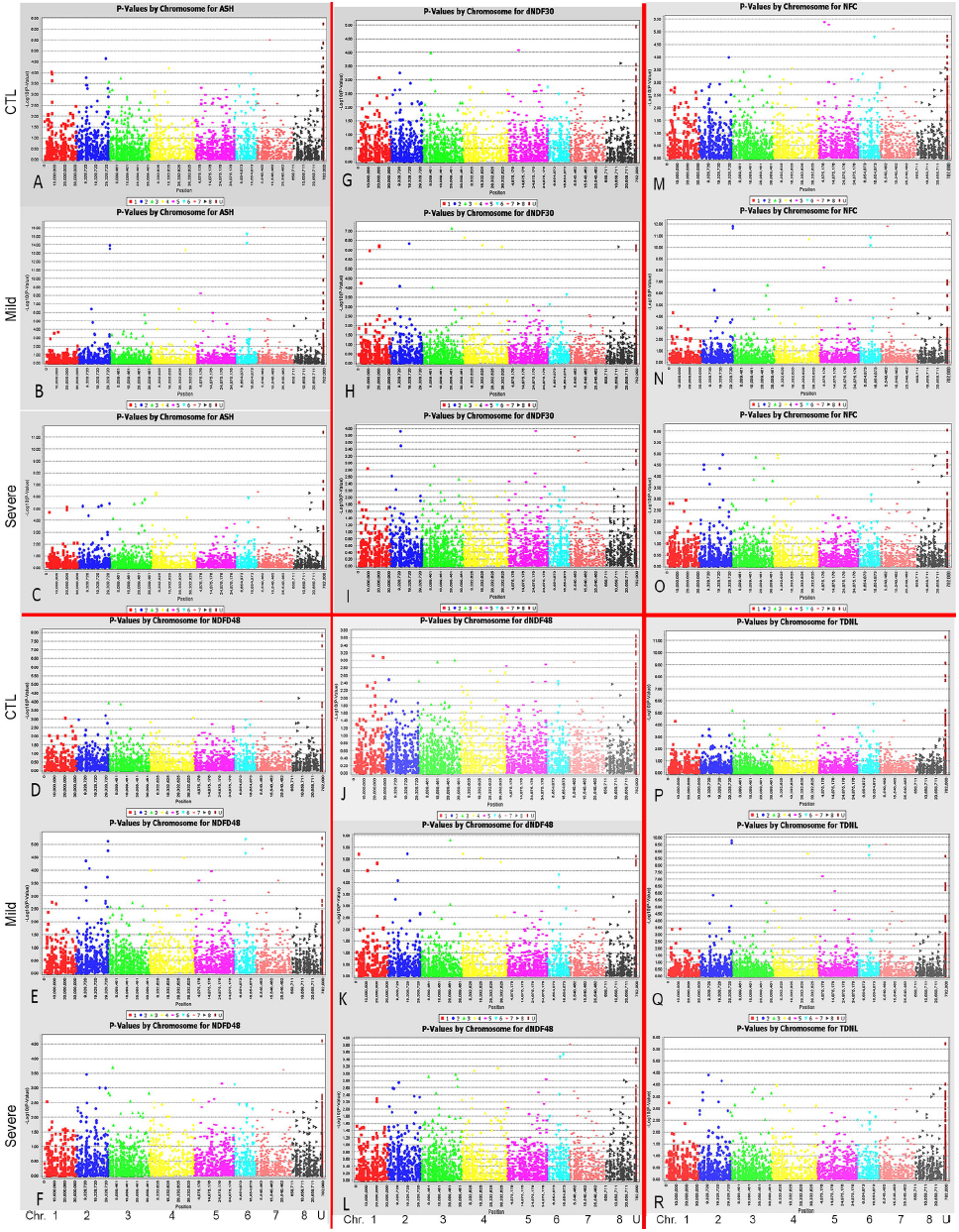
Manhattan plots of marker-trait association of six most significant quality traits under well-watered control (CTL), mild water deficit (Mild) and severe water deficit (Severe). The X-axis presents chromosome positions of loci based on the reference genome of *M. truncatula* (Mt4.0, v1). The Y-axis shows negative log (P-values) of marker-trait association. Chromosome numbers were assigned and illustrated at the bottom of the figures.

When phenotypic means were used for marker-trait association, similar association profile was found among DRIMI, DMI1, RFV, and RFQ (Fig. 7.1 A-D), and most significant markers were identified on chromosome 8 and unknown chromosome (Chr. U). Similar profiles were also found among ADF, DDM, TDN, ENE, ME, NEM, NEG and NEL (Fig. 7.1 E-L) and most significant markers were located on chromosome 8 and U. The same was true between DM and moisture (Fig. 7.1 M and N). Although similar association patterns were found between in vitro dry matter digestibility 30h and 48h (IV30 and IV48, respectively), none was statistically significant, neither in IV30 nor in IV48 (Fig. 7.1 O and P). Different association profiles were shown among the remaining traits (Fig. 7.2 A-O). Among those, significant markers were found on chromosomes 1, 2, 3 and 8 for dNDF48 (Fig. 7.2 C), whereas no significant marker was identified in dNDF30 (Fig. 7.2 B). One significant marker was identified in dNDF48 but its chromosome location was unknown (Fig. 7.2 D). Significant markers were identified on chromosomes 3 and 5 for lignin content (Fig. 7.2 E). A marker was identified for NFC, with unknown chromosome position (Fig. 7.2 F). Seven markers were highly significantly associated with ash and they were located on chromosomes 1, 2, 3, 7 and 8 (Fig. 7.2 J). Only one marker on chromosome 4 was significantly associated with fat (Fig. 7.2 K).

**Fig. 7.1.**
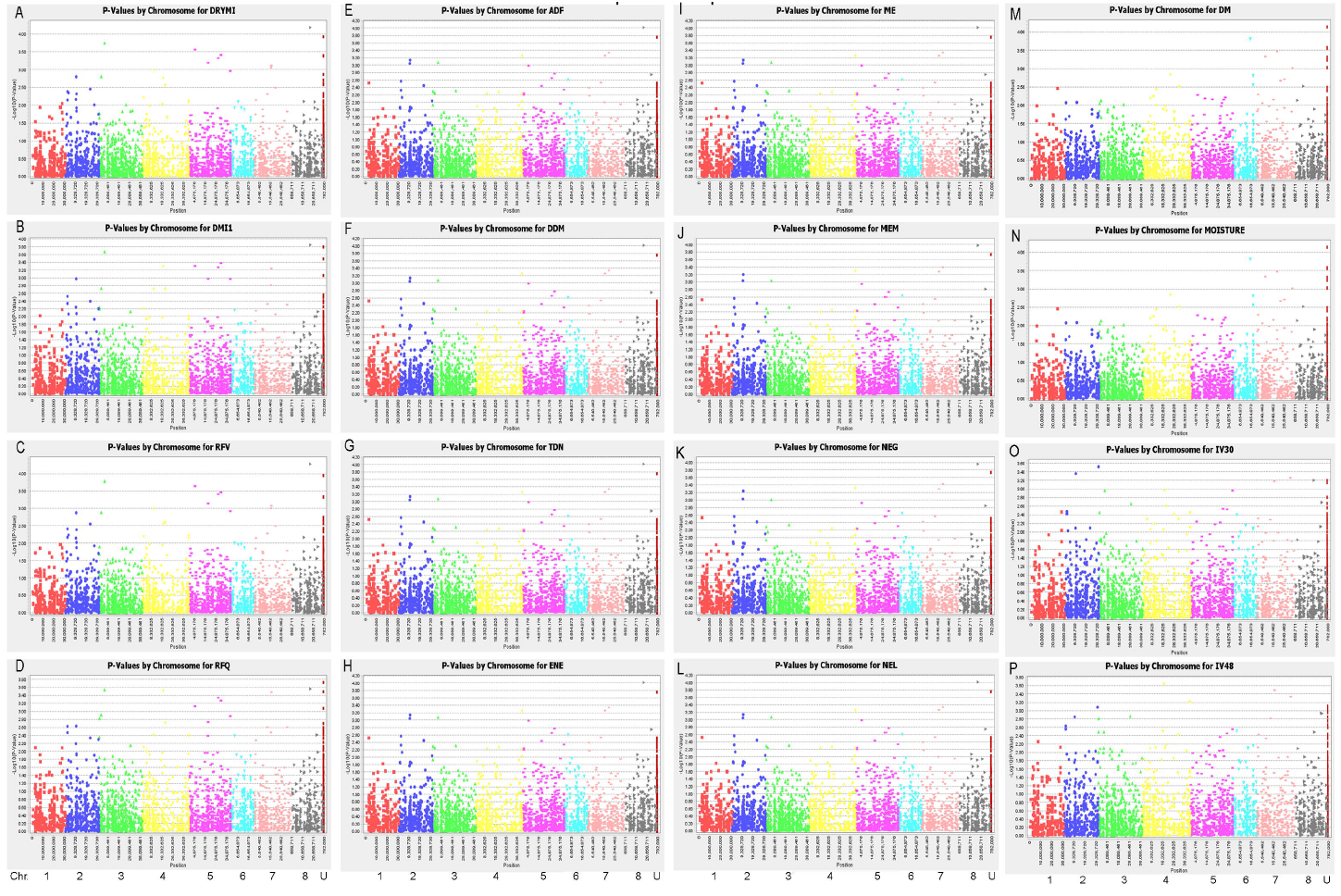
Comparison of association profiles of forage quality traits using means of three irrigation treatments. The X-axis presents chromosome positions of loci based on the reference genome of *M. truncatula* (Mt4.0, v1). The Y-axis shows negative log (P-values) of marker-trait association. Chromosome numbers are illustrated by colors at the bottom of the figures.

**Fig. 7.2.**
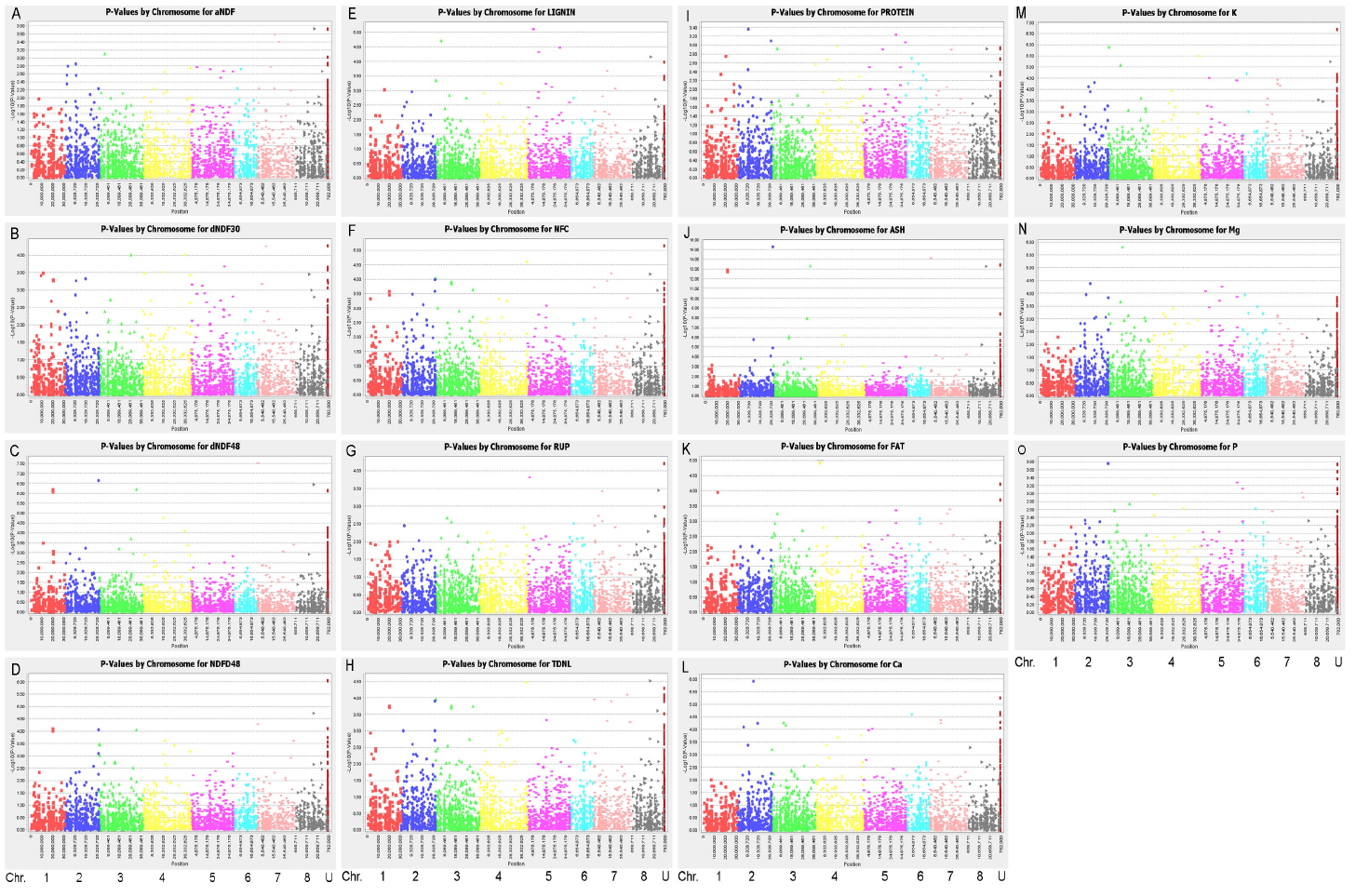
Comparison of association profiles of forage quality traits using means of three irrigation treatments. The X-axis presents chromosome positions of loci based on the reference genome of *M. truncatula* (Mt4.0, v1). The Y-axis shows negative log (P-values) of marker-trait association. Chromosome numbers are illustrated by colors at the bottom of the figures.

### Common markers identified among multiple quality traits and different treatments

Despite different loci identified among quality traits, common marker loci were found among multiple traits (Table 3). Marker S1_110050725 on chromosome 4 identified in CTL for ADF also significantly associated with other 10 traits including DDM, ENE, IVDMD30, IVDMD48, ME, NEG, NEL NEM, Protein and TDN (Table 3, top row). Similarly, markers S1_305729816 for DMI1 in CTL also associated with 6 other traits: IVDMD30, IVDMD48, NDFD48, RFQ, TNDL and protein. Marker S1_231443201 for ADF shared its association with DDM, ENE, ME, NEG, NEL, NEM, TDN and TDNL. Four markers (S1_197238737 - 90) at the same locus on chromosome 6 and unknown marker S1_292679040 identified for ash in the mild stress were also associated with 6 other traits. Markers S1_351118210 and S1_276968305 identified in CTL for IVDMD48 and ash, respectively, were significantly associated with 5 other traits. Marker S1_174013573 identified in the severe stress for DMI1 were also associated with DRYMI, fat, RFQ and RFV. There were eight markers associated ash were also associated with NDFD48, NFC and TDNL (Table 3). There were nine, eighteen and fifty-five markers were shared their associations with three, two and one traits, respectively. The remaining markers were independently associated with one trait (Table 3, bottom part). Most of the high significant markers with lower P-value and higher R^2^ are among of common markers, suggesting these markers may have major effects on the respective traits. Among those, ten markers were associated with the same three traits (NDFD48, NFC and TDN). The P-values of these markers ranged from 4.01E-08 (S1_21394491) to 5.79E-16 (S1_197238737) and the marker’s R^2^ ranged from 0.22 to 0.38, respectively (Table 3).

**Table 3.**
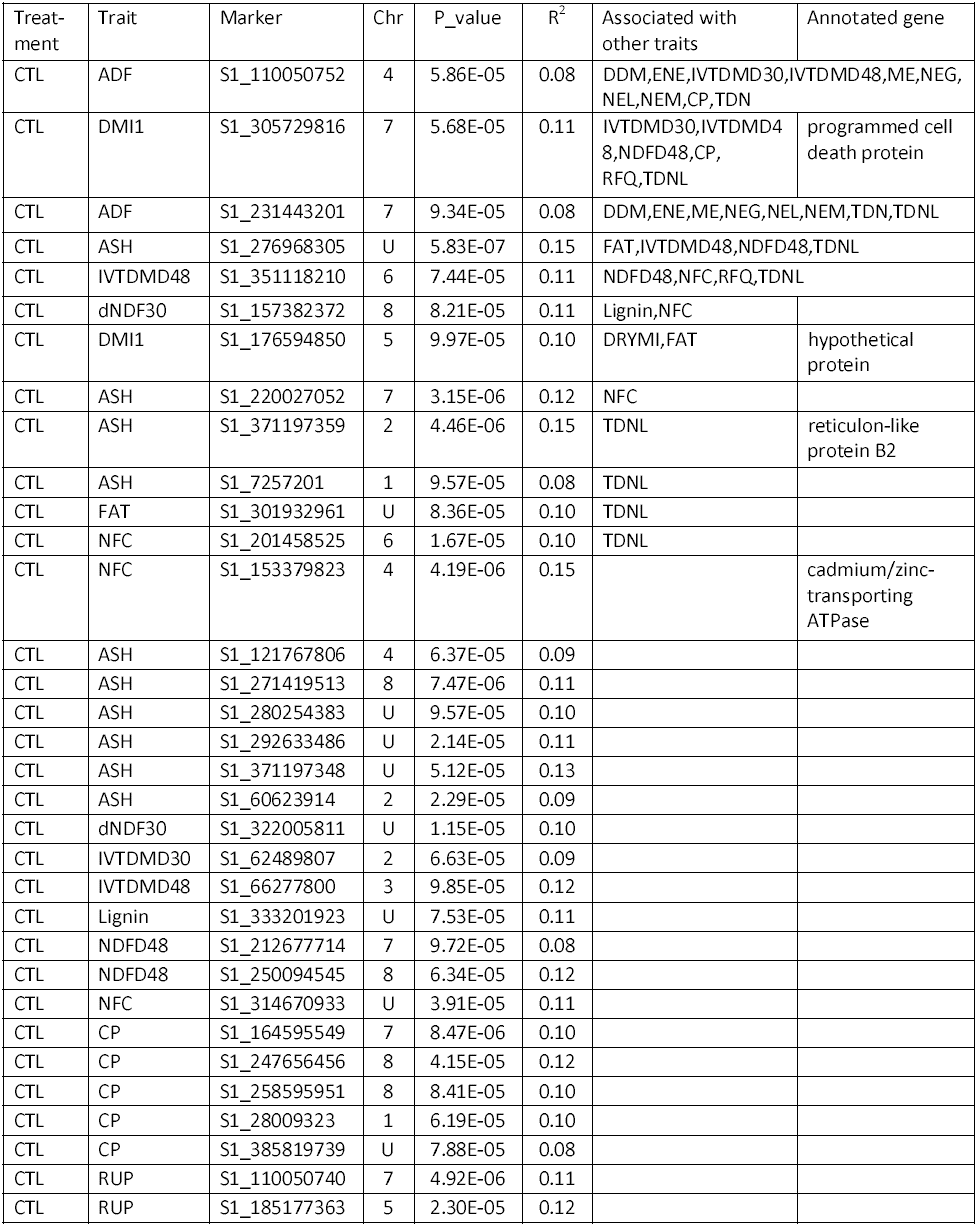
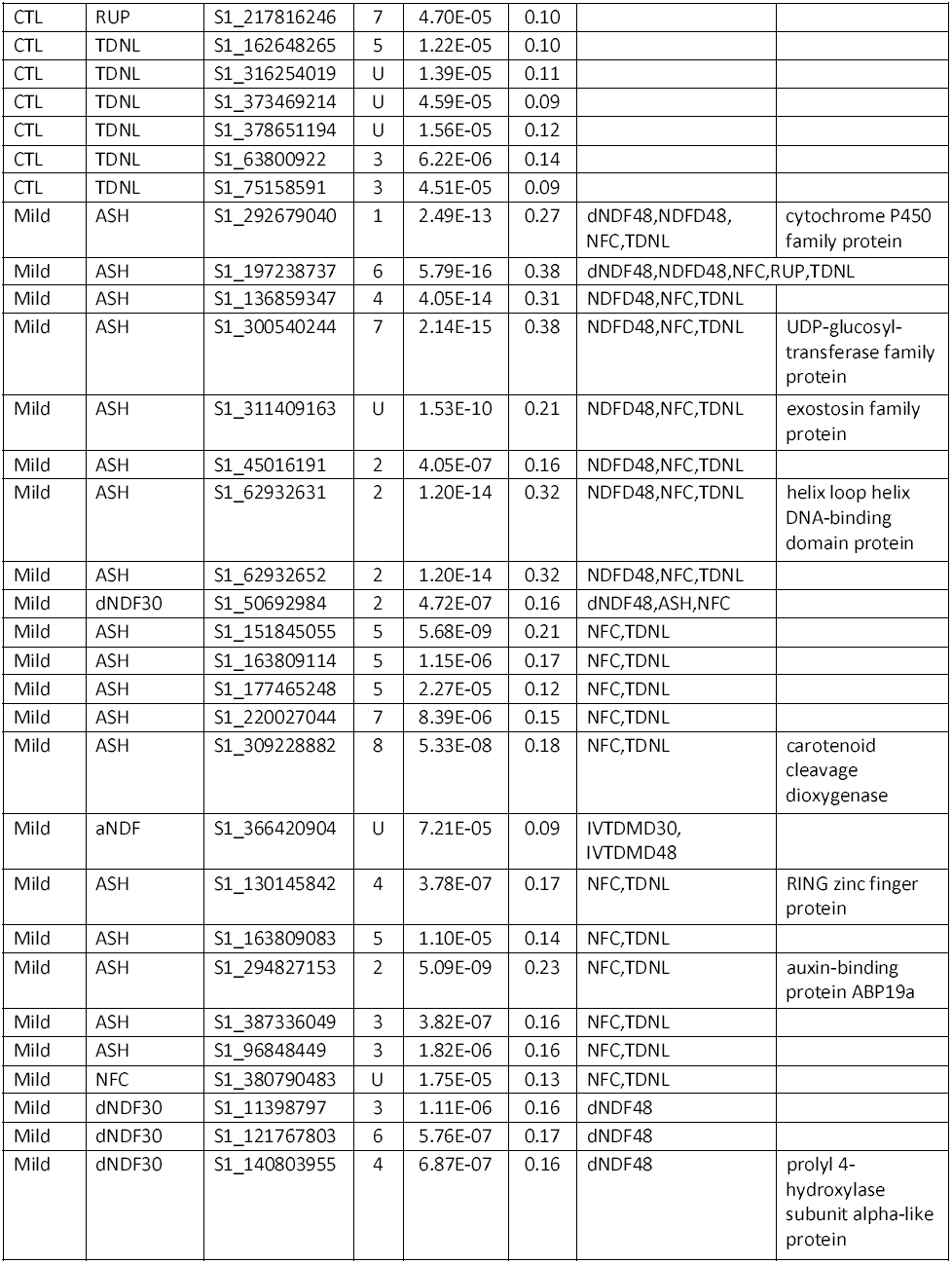
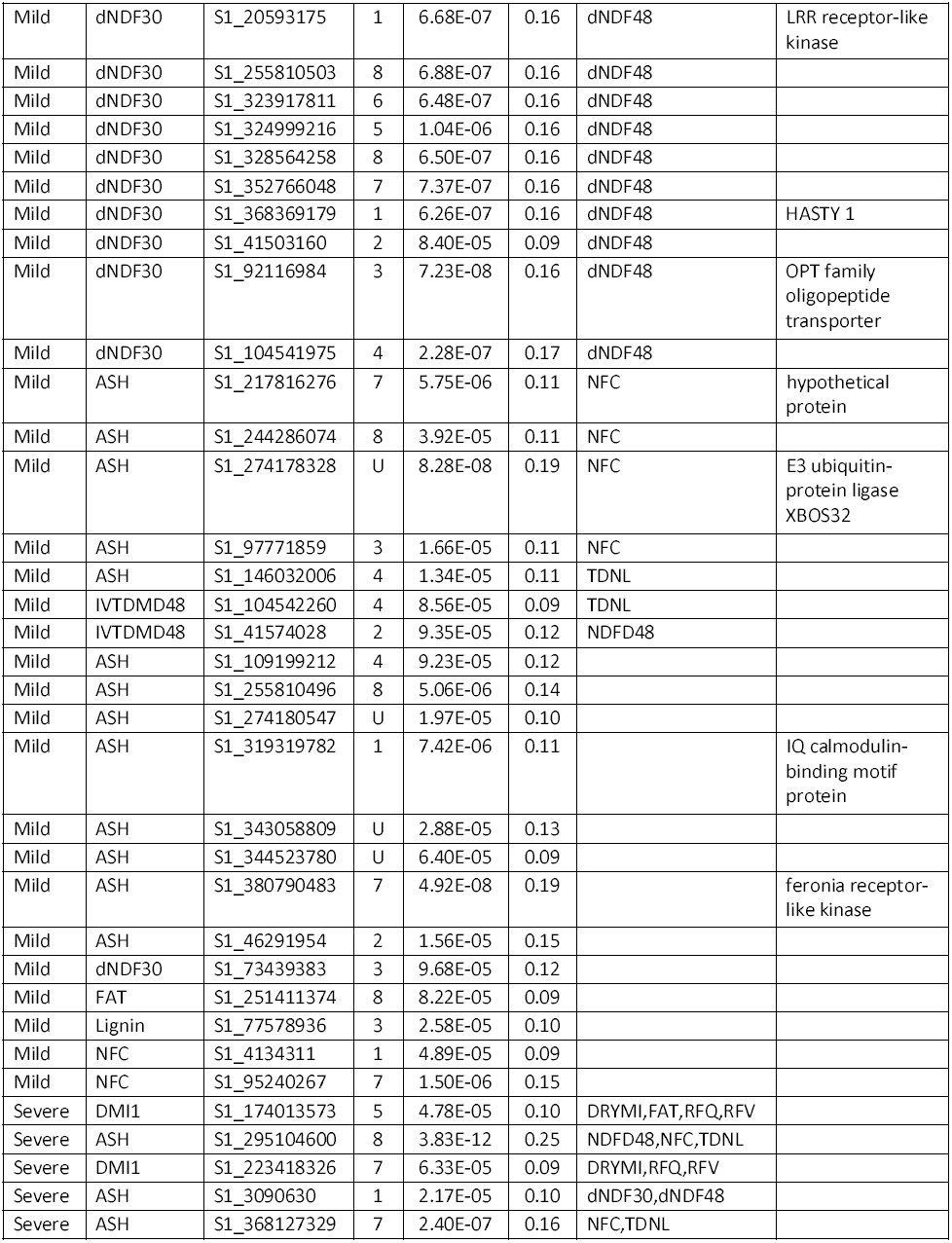
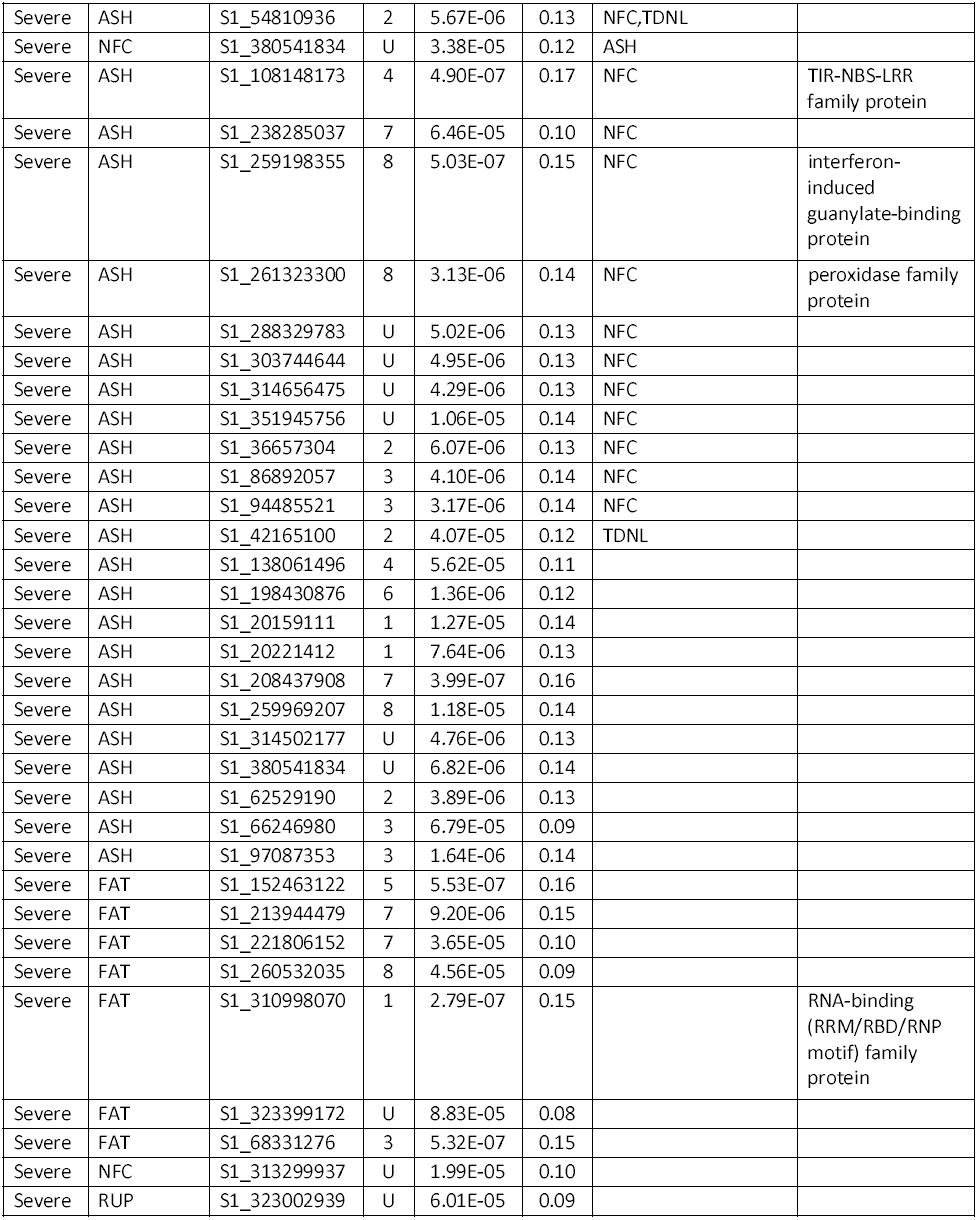
Significant markers associated with forage quality traits under well-watered control (CTL), mild water deficit (Mild) and severe water deficit (Severe) in the panel of 198 accessions

| Treat-<br>ment | Trait | Marker | Chr | P_value | R<sup>2</sup> | Associated with<br>other traits | Annotated gene |
| --- | --- | --- | --- | --- | --- | --- | --- |
| CTL | ADF | S1_110050752 | 4 | 5.86E-05 | 0.08 | DDM,ENE,IVTDMD30,IVTDMD48,ME,NEG,<br>NEL,NEM,CP,TDN |  |
| CTL | DMI1 | S1_305729816 | 7 | 5.68E-05 | 0.11 | IVTDMD30,IVTDMD4<br>8,NDFD48,CP,<br>RFQ,TDNL | programmed cell<br>death protein |
| CTL | ADF | S1_231443201 | 7 | 9.34E-05 | 0.08 | DDM,ENE,ME,NEG,NEL,NEM,TDN,TDNL |  |
| CTL | ASH | S1_276968305 | U | 5.83E-07 | 0.15 | FAT,IVTDMD48,NDFD48,TDNL |  |
| CTL | IVTDMD48 | S1_351118210 | 6 | 7.44E-05 | 0.11 | NDFD48,NFC,RFQ,TDNL |  |
| CTL | dNDF30 | S1_157382372 | 8 | 8.21E-05 | 0.11 | Lignin,NFC |  |
| CTL | DMI1 | S1_176594850 | 5 | 9.97E-05 | 0.10 | DRYMI,FAT | hypothetical<br>protein |
| CTL | ASH | S1_220027052 | 7 | 3.15E-06 | 0.12 | NFC |  |
| CTL | ASH | S1_371197359 | 2 | 4.46E-06 | 0.15 | TDNL | reticulon-like<br>protein B2 |
| CTL | ASH | S1_7257201 | 1 | 9.57E-05 | 0.08 | TDNL |  |
| CTL | FAT | S1_301932961 | U | 8.36E-05 | 0.10 | TDNL |  |
| CTL | NFC | S1_201458525 | 6 | 1.67E-05 | 0.10 | TDNL |  |
| CTL | NFC | S1_153379823 | 4 | 4.19E-06 | 0.15 |  | cadmium/zinc-<br>transporting<br>ATPase |
| CTL | ASH | S1_121767806 | 4 | 6.37E-05 | 0.09 |  |  |
| CTL | ASH | S1_271419513 | 8 | 7.47E-06 | 0.11 |  |  |
| CTL | ASH | S1_280254383 | U | 9.57E-05 | 0.10 |  |  |
| CTL | ASH | S1_292633486 | U | 2.14E-05 | 0.11 |  |  |
| CTL | ASH | S1_371197348 | U | 5.12E-05 | 0.13 |  |  |
| CTL | ASH | S1_60623914 | 2 | 2.29E-05 | 0.09 |  |  |
| CTL | dNDF30 | S1_322005811 | U | 1.15E-05 | 0.10 |  |  |
| CTL | IVTDMD30 | S1_62489807 | 2 | 6.63E-05 | 0.09 |  |  |
| CTL | IVTDMD48 | S1_66277800 | 3 | 9.85E-05 | 0.12 |  |  |
| CTL | Lignin | S1_333201923 | U | 7.53E-05 | 0.11 |  |  |
| CTL | NDFD48 | S1_212677714 | 7 | 9.72E-05 | 0.08 |  |  |
| CTL | NDFD48 | S1_250094545 | 8 | 6.34E-05 | 0.12 |  |  |
| CTL | NFC | S1_314670933 | U | 3.91E-05 | 0.11 |  |  |
| CTL | CP | S1_164595549 | 7 | 8.47E-06 | 0.10 |  |  |
| CTL | CP | S1_247656456 | 8 | 4.15E-05 | 0.12 |  |  |
| CTL | CP | S1_258595951 | 8 | 8.41E-05 | 0.10 |  |  |
| CTL | CP | S1_28009323 | 1 | 6.19E-05 | 0.10 |  |  |
| CTL | CP | S1_385819739 | U | 7.88E-05 | 0.08 |  |  |
| CTL | RUP | S1_110050740 | 7 | 4.92E-06 | 0.11 |  |  |
| CTL | RUP | S1_185177363 | 5 | 2.30E-05 | 0.12 |  |  |
| CTL | RUP | S1_217816246 | 7 | 4.70E-05 | 0.10 |  |  |
| CTL | TDNL | S1_162648265 | 5 | 1.22E-05 | 0.10 |  |  |
| CTL | TDNL | S1_316254019 | U | 1.39E-05 | 0.11 |  |  |
| CTL | TDNL | S1_373469214 | U | 4.59E-05 | 0.09 |  |  |
| CTL | TDNL | S1_378651194 | U | 1.56E-05 | 0.12 |  |  |
| CTL | TDNL | S1_63800922 | 3 | 6.22E-06 | 0.14 |  |  |
| CTL | TDNL | S1_75158591 | 3 | 4.51E-05 | 0.09 |  |  |
| Mild | ASH | S1_292679040 | 1 | 2.49E-13 | 0.27 | dNDF48,NDFD48,<br>NFC,TDNL | cytochrome P450<br>family protein |
| Mild | ASH | S1_197238737 | 6 | 5.79E-16 | 0.38 | dNDF48,NDFD48,NFC,RUP,TDNL |  |
| Mild | ASH | S1_136859347 | 4 | 4.05E-14 | 0.31 | NDFD48,NFC,TDNL |  |
| Mild | ASH | S1_300540244 | 7 | 2.14E-15 | 0.38 | NDFD48,NFC,TDNL | UDP-glucosyl-<br>transferase family<br>protein |
| Mild | ASH | S1_311409163 | U | 1.53E-10 | 0.21 | NDFD48,NFC,TDNL | exostosin family<br>protein |
| Mild | ASH | S1_45016191 | 2 | 4.05E-07 | 0.16 | NDFD48,NFC,TDNL |  |
| Mild | ASH | S1_62932631 | 2 | 1.20E-14 | 0.32 | NDFD48,NFC,TDNL | helix loop helix<br>DNA-binding<br>domain protein |
| Mild | ASH | S1_62932652 | 2 | 1.20E-14 | 0.32 | NDFD48,NFC,TDNL |  |
| Mild | dNDF30 | S1_50692984 | 2 | 4.72E-07 | 0.16 | dNDF48,ASH,NFC |  |
| Mild | ASH | S1_151845055 | 5 | 5.68E-09 | 0.21 | NFC,TDNL |  |
| Mild | ASH | S1_163809114 | 5 | 1.15E-06 | 0.17 | NFC,TDNL |  |
| Mild | ASH | S1_177465248 | 5 | 2.27E-05 | 0.12 | NFC,TDNL |  |
| Mild | ASH | S1_220027044 | 7 | 8.39E-06 | 0.15 | NFC,TDNL |  |
| Mild | ASH | S1_309228882 | 8 | 5.33E-08 | 0.18 | NFC,TDNL | carotenoid<br>cleavage<br>dioxygenase |
| Mild | aNDF | S1_366420904 | U | 7.21E-05 | 0.09 | IVTDM30,<br>IVTDM48 |  |
| Mild | ASH | S1_130145842 | 4 | 3.78E-07 | 0.17 | NFC,TDNL | RING zinc finger<br>protein |
| Mild | ASH | S1_163809083 | 5 | 1.10E-05 | 0.14 | NFC,TDNL |  |
| Mild | ASH | S1_294827153 | 2 | 5.09E-09 | 0.23 | NFC,TDNL | auxin-binding<br>protein ABP19a |
| Mild | ASH | S1_387336049 | 3 | 3.82E-07 | 0.16 | NFC,TDNL |  |
| Mild | ASH | S1_96848449 | 3 | 1.82E-06 | 0.16 | NFC,TDNL |  |
| Mild | NFC | S1_380790483 | U | 1.75E-05 | 0.13 | NFC,TDNL |  |
| Mild | dNDF30 | S1_11398797 | 3 | 1.11E-06 | 0.16 | dNDF48 |  |
| Mild | dNDF30 | S1_121767803 | 6 | 5.76E-07 | 0.17 | dNDF48 |  |
| Mild | dNDF30 | S1_140803955 | 4 | 6.87E-07 | 0.16 | dNDF48 | prolyl 4-hydroxylase<br>subunit alpha-like<br>protein |
| Mild | dNDF30 | S1_20593175 | 1 | 6.68E-07 | 0.16 | dNDF48 | LRR receptor-like kinase |
| Mild | dNDF30 | S1_255810503 | 8 | 6.88E-07 | 0.16 | dNDF48 |  |
| Mild | dNDF30 | S1_323917811 | 6 | 6.48E-07 | 0.16 | dNDF48 |  |
| Mild | dNDF30 | S1_324999216 | 5 | 1.04E-06 | 0.16 | dNDF48 |  |
| Mild | dNDF30 | S1_328564258 | 8 | 6.50E-07 | 0.16 | dNDF48 |  |
| Mild | dNDF30 | S1_352766048 | 7 | 7.37E-07 | 0.16 | dNDF48 |  |
| Mild | dNDF30 | S1_368369179 | 1 | 6.26E-07 | 0.16 | dNDF48 | HASTY 1 |
| Mild | dNDF30 | S1_41503160 | 2 | 8.40E-05 | 0.09 | dNDF48 |  |
| Mild | dNDF30 | S1_92116984 | 3 | 7.23E-08 | 0.16 | dNDF48 | OPT family oligopeptide transporter |
| Mild | dNDF30 | S1_104541975 | 4 | 2.28E-07 | 0.17 | dNDF48 |  |
| Mild | ASH | S1_217816276 | 7 | 5.75E-06 | 0.11 | NFC | hypothetical protein |
| Mild | ASH | S1_244286074 | 8 | 3.92E-05 | 0.11 | NFC |  |
| Mild | ASH | S1_274178328 | U | 8.28E-08 | 0.19 | NFC | E3 ubiquitin-protein ligase XBOS32 |
| Mild | ASH | S1_97771859 | 3 | 1.66E-05 | 0.11 | NFC |  |
| Mild | ASH | S1_146032006 | 4 | 1.34E-05 | 0.11 | TDNL |  |
| Mild | IVTDM48 | S1_104542260 | 4 | 8.56E-05 | 0.09 | TDNL |  |
| Mild | IVTDM48 | S1_41574028 | 2 | 9.35E-05 | 0.12 | NDFD48 |  |
| Mild | ASH | S1_109199212 | 4 | 9.23E-05 | 0.12 |  |  |
| Mild | ASH | S1_255810496 | 8 | 5.06E-06 | 0.14 |  |  |
| Mild | ASH | S1_274180547 | U | 1.97E-05 | 0.10 |  |  |
| Mild | ASH | S1_319319782 | 1 | 7.42E-06 | 0.11 |  | IQ calmodulin-binding motif protein |
| Mild | ASH | S1_343058809 | U | 2.88E-05 | 0.13 |  |  |
| Mild | ASH | S1_344523780 | U | 6.40E-05 | 0.09 |  |  |
| Mild | ASH | S1_380790483 | 7 | 4.92E-08 | 0.19 |  | feronia receptor-like kinase |
| Mild | ASH | S1_46291954 | 2 | 1.56E-05 | 0.15 |  |  |
| Mild | dNDF30 | S1_73439383 | 3 | 9.68E-05 | 0.12 |  |  |
| Mild | FAT | S1_251411374 | 8 | 8.22E-05 | 0.09 |  |  |
| Mild | Lignin | S1_77578936 | 3 | 2.58E-05 | 0.10 |  |  |
| Mild | NFC | S1_4134311 | 1 | 4.89E-05 | 0.09 |  |  |
| Mild | NFC | S1_95240267 | 7 | 1.50E-06 | 0.15 |  |  |
| Severe | DMI1 | S1_174013573 | 5 | 4.78E-05 | 0.10 | DRYMI,FAT,RFQ,RFV |  |
| Severe | ASH | S1_295104600 | 8 | 3.83E-12 | 0.25 | NDFD48,NFC,TDNL |  |
| Severe | DMI1 | S1_223418326 | 7 | 6.33E-05 | 0.09 | DRYMI,RFQ,RFV |  |
| Severe | ASH | S1_3090630 | 1 | 2.17E-05 | 0.10 | dNDF30,dNDF48 |  |
| Severe | ASH | S1_368127329 | 7 | 2.40E-07 | 0.16 | NFC,TDNL |  |
| Severe | ASH | S1_54810936 | 2 | 5.67E-06 | 0.13 | NFC,TDNL |  |
| Severe | NFC | S1_380541834 | U | 3.38E-05 | 0.12 | ASH |  |
| Severe | ASH | S1_108148173 | 4 | 4.90E-07 | 0.17 | NFC | TIR-NBS-LRR family protein |
| Severe | ASH | S1_238285037 | 7 | 6.46E-05 | 0.10 | NFC |  |
| Severe | ASH | S1_259198355 | 8 | 5.03E-07 | 0.15 | NFC | interferon-induced guanylate-binding protein |
| Severe | ASH | S1_261323300 | 8 | 3.13E-06 | 0.14 | NFC | peroxidase family protein |
| Severe | ASH | S1_288329783 | U | 5.02E-06 | 0.13 | NFC |  |
| Severe | ASH | S1_303744644 | U | 4.95E-06 | 0.13 | NFC |  |
| Severe | ASH | S1_314656475 | U | 4.29E-06 | 0.13 | NFC |  |
| Severe | ASH | S1_351945756 | U | 1.06E-05 | 0.14 | NFC |  |
| Severe | ASH | S1_36657304 | 2 | 6.07E-06 | 0.13 | NFC |  |
| Severe | ASH | S1_86892057 | 3 | 4.10E-06 | 0.14 | NFC |  |
| Severe | ASH | S1_94485521 | 3 | 3.17E-06 | 0.14 | NFC |  |
| Severe | ASH | S1_42165100 | 2 | 4.07E-05 | 0.12 | TDNL |  |
| Severe | ASH | S1_138061496 | 4 | 5.62E-05 | 0.11 |  |  |
| Severe | ASH | S1_198430876 | 6 | 1.36E-06 | 0.12 |  |  |
| Severe | ASH | S1_20159111 | 1 | 1.27E-05 | 0.14 |  |  |
| Severe | ASH | S1_20221412 | 1 | 7.64E-06 | 0.13 |  |  |
| Severe | ASH | S1_208437908 | 7 | 3.99E-07 | 0.16 |  |  |
| Severe | ASH | S1_259969207 | 8 | 1.18E-05 | 0.14 |  |  |
| Severe | ASH | S1_314502177 | U | 4.76E-06 | 0.13 |  |  |
| Severe | ASH | S1_380541834 | U | 6.82E-06 | 0.14 |  |  |
| Severe | ASH | S1_62529190 | 2 | 3.89E-06 | 0.13 |  |  |
| Severe | ASH | S1_66246980 | 3 | 6.79E-05 | 0.09 |  |  |
| Severe | ASH | S1_97087353 | 3 | 1.64E-06 | 0.14 |  |  |
| Severe | FAT | S1_152463122 | 5 | 5.53E-07 | 0.16 |  |  |
| Severe | FAT | S1_213944479 | 7 | 9.20E-06 | 0.15 |  |  |
| Severe | FAT | S1_221806152 | 7 | 3.65E-05 | 0.10 |  |  |
| Severe | FAT | S1_260532035 | 8 | 4.56E-05 | 0.09 |  |  |
| Severe | FAT | S1_310998070 | 1 | 2.79E-07 | 0.15 |  | RNA-binding (RRM/RBD/RNP motif) family protein |
| Severe | FAT | S1_323399172 | U | 8.83E-05 | 0.08 |  |  |
| Severe | FAT | S1_68331276 | 3 | 5.32E-07 | 0.15 |  |  |
| Severe | NFC | S1_313299937 | U | 1.99E-05 | 0.10 |  |  |
| Severe | RUP | S1_323002939 | U | 6.01E-05 | 0.09 |  |  |

To oversee the genetic architecture of the population under different treatments, we compared markers significantly associated with CTL, mild and severe drought treatments with those identified using mean values of all treatments. Among 68 markers identified in the control, 17 were also identified in the mean (Fig. 8 A). Of 70 significant markers identified in the mild stress, only 10 were also identified in the mean (Fig. 8 A). Among 67 markers identified in severe drought, 20 were also found in the mean (Fig. 8 A). We have also compared the common markers identified among the three treatments directly. There were 3 common markers between each pair of treatments (Fig. 8 B). Only 2 markers were found in all three treatments (Fig. 8 B).

**Fig. 8.**
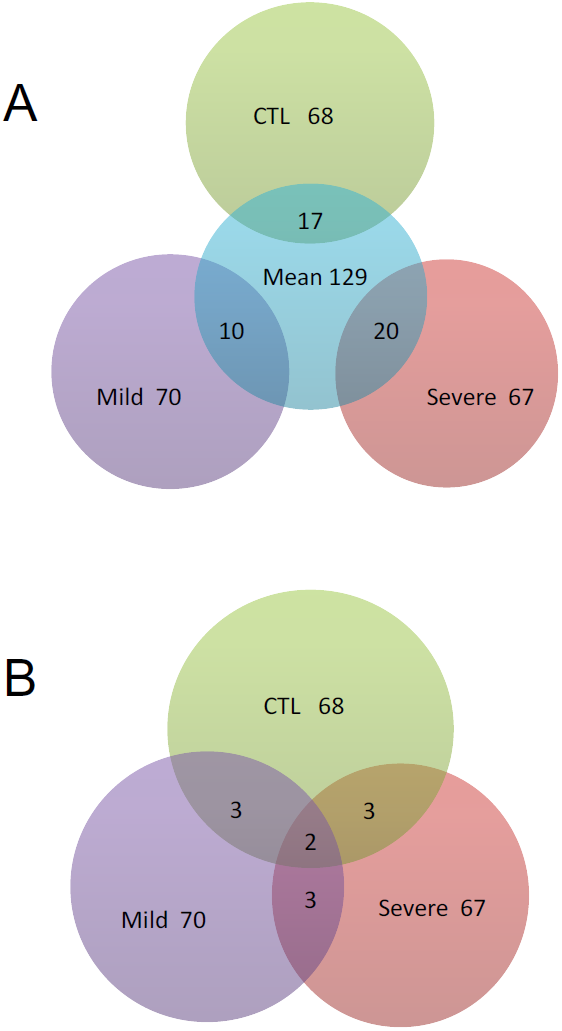
A Vann chart of significant loci associated with forage quality resulting from GWAS for quality traits in alfalfa under well-watered (CTL), mild and severe water deficits compared with mean (A) and without mean (B). The numbers of significant loci identified under each treatment were compared with those of mean values of all treatments, showing the numbers of common (overlapped) and specific loci for different treatments.

### Assignment of significant markers to annotated genes

Using the flanking sequences of the significant markers, we performed a BLAST search to identify potential candidate genes linked to significant marker loci. Of significant markers identified, 23 markers were found to be linked to known genes in the *M. truncatula* genome Mtv4.2 (Table3). Of those identified under well-watered condition (CTL), marker S1_305729816 and S1_176594850 associated with DMI1 were linked to a programmed cell death and hypothetic proteins, respectively. Marker S1_371197359 associated with ash was linked to a reticulon-like protein B2. Two markers (S1_153379823 and S1_ 153379823) located at the same locus associated with NFC was linked to cadmium/zinc transporting ATPase. Of those identified under mild water deficit, many markers were associated with ash. Among them, marker S1_292679040 was linked to cytochrome P450 family protein; marker S1_300540244 was linked to UDP-glucosyltransferase; S1_311409163 was linked to exostosin; S1_62932631 was linked to helix loop helix DNA-binding domain protein; S1_309228882 was linked to carotenoid cleavage dioxygenase; S1_130145842 was linked to RING zinc finger protein; S1_294827153 was linked to auxin-binding protein ABP19a; S1_274178328 was linked to E3 ubiquitin protein ligase XBOS32; S1_319319782 was linked to IQ calmodulin-binding motif protein; and S1_380790483 was linked to Feronia receptor-like kinase. Three markers, S1_20593175, S1_368369179 and S1_92116984 associated with dNDF30 and dNDF48 were linked to LRR-receptor-like kinase, HASTY1 and OPT family oligopeptide transporter, respectively. Of those identified under severe water deficit, markers S1_108148173, S1_259198355 and S1_261323300 were linked to TIR-NBS-LRR resistance protein, interferon-induced guanylate-binding protein and peroxidase family protein, respectively. Anther marker S1_310998070 associated with fat was linked to RNA-binding (RRM/RBD/RNP motif) protein.

## Discussion

### Phenotypic variations of forage quality among alfalfa accessions

Near Infrared Reflectance Spectroscopy (NIRS) has become popular for use in measuring forage quality factors, since relatively precise results can be obtained fast with a proper NIRS. In the present report, we used NIRS results for 26 forage quality factors including protein, fibers, energy and minerals and performed cluster analysis on phenotypic variations of forage quality among 198 alfalfa accessions. A majority accessions were classified into the higher quality cluster, among those, two drought sensitive varieties were in this cluster. The remaining accessions were grouped into the lower quality cluster where three drought tolerance varieties were in this cluster. There is a trend that drought sensitive varieties have relative higher quality than those with drought tolerance. It suggests that breeding for drought tolerant alfalfa may result quality penalty as the crop diverts energy to implementing drought resistance. Therefore, quality factors should be considered when breeding for abiotic tolerance.

Commercial markets are seeking common means for estimating forage quality in terms of its value as a feed for livestock. Currently, RFV is the most popular quality measurement and is accepted by hay buyers. Dairy’s often retest hay and look for NEL, CP and aNDF are excellent indicators of energy, protein and fiber, respectively. In this report, water deficit significantly decreased aNDF and slightly decreased CP but increased NEL (Fig. 1). Relative Feed Value is calculated from ADF an old estimate of digestible dry matter and NDF and estimate of dry matter intake that is still used (Undersander and Moore, 2004). In the present study, we found a high positive correlation (R^2^=9.405) between RFV and the mean of 26 quality factors (Fig. 9 A). However, the RFV is not a good measurement of NDF digestibility. The RFQ index has been developed to overcome this shortage as it takes into consideration the differences in digestibility of the fiber fraction and can be used to more accurately predict animal performance. We therefore performed correlation analysis between RFQ and the mean of 26quality factors and obtained very high correlation coefficient (R^2^=0.9946) between them (Fig. 9B). Our results support that the RFQ is more accurate in estimating overall forage quality compared to the RFV. Differences in the digestibility of the fiber fraction can result in a difference in animal performance even if the forages with the same RFV are fed. The RFQ index is an improvement over the RFV index for estimating forage quality as it better reflects the performance of animal fed the forage.

**Fig. 9.**
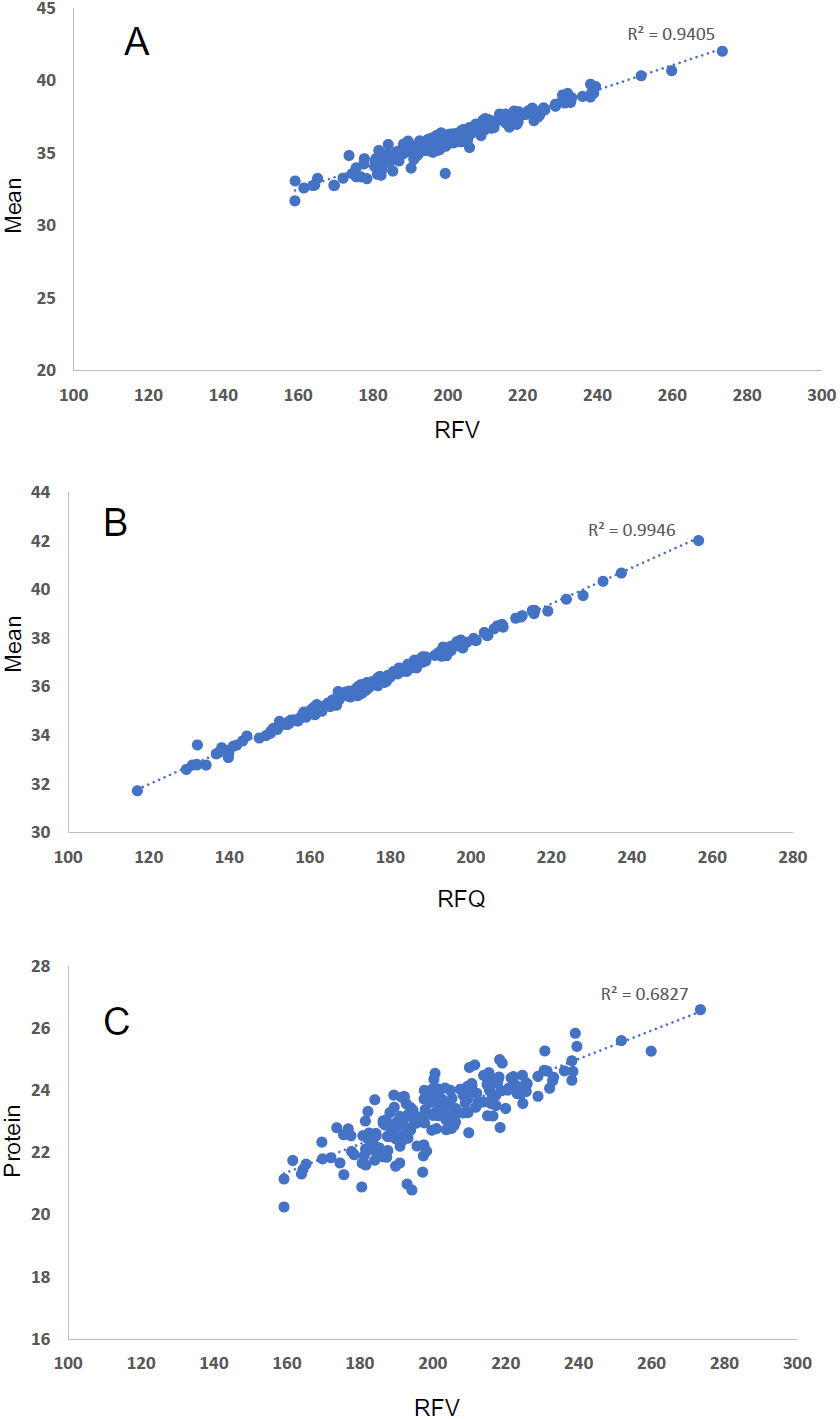
Correlation coefficients between means of quality traits and RFV (A) or RFQ (B) and between RFV and crude protein content (C).

On the other hand, neither RFV nor RFQ does not include protein concentration or physical characteristics. Protein and physical characteristics are important factors for evaluating forage quality. To understand the relationships between those factors, in the present study, we analyzed the correlations between protein content and the RFV (Fig. 9 C). Although positive correlations were found between protein and the RFV, the correlation was moderate (R^2^=0.6827) between them. Similar correlation coefficient (R^2^=0.6447) was also found between the RFQ and the protein content (data not shown). Therefore, protein and physical characteristics should be evaluated along with RFV and RFQ for a complete assessment of forage quality.

### Mild drought intends to decrease fiber content and improve digestibility in alfalfa

Production of alfalfa for maximum yield and quality requires an understanding of how environmental and cultural factors influence crop growth and development. These factors include growing conditions, stage of maturity at harvest, and climate conditions. Causations exists between the environment, plant response, and nutritive value. In general, yield and forage quality are inversely related. Any factor that retards plant development tends to promote the maintenance of forage quality. If a plant is stressed during growth, a shorter, finer-stemmed, leafier alfalfa is often produced. On the other hand, factors that accelerate growth, such as drought and high temperature, tend to have a negative impact on forage quality. Alfalfa is relatively drought tolerant because its deeper root systems allow alfalfa to absorb deep soil water and quickly recover from drought conditions. However, when transpiration exceeds water absorption, a stress is imposed on the plant influencing metabolism, development, growth, and ultimately yield. Water deficit promotes a reduction in vegetative growth and promote early maturity. It has been suggested that mild drought stress may be beneficial for forage quality as alfalfa as drought-stressed alfalfa will accelerate its shift to reproductive growth (Cassida, 2012). Once a stem has flowered, there will be little additional growth in the stem. Stem internode growth is suppressed, resulting in a greater proportion of leaves on shorter stems. In a short term of drought stress, the greater proportion of leaves in the drought-stressed forage improves feed quality and digestibility. However, if drought stress has been too severe, and for an extended period, plant stress is permanent and may not be recovered.

Alfalfa fiber is consisted of three components: cellulose, hemicellulose and lignin. Increasing fiber content of a forage generally decreases its energy content. Of the fiber fractions, cellulose is the major compound digested by the animal while lignin is virtually indigestible in both the rumen and lower intestines. In our study, drought decreased significantly both ADF and NDF, which in turn increased energy-related traits such as TDN, ENE, DDM, NFC, RFV and RFQ (Fig. 1). Drought affects forage quality as cell wall remodeling is a common plant response to abiotic stresses. Biomass composition was drastically altered due to drought stress, with significant reductions in cell wall and cellulose content. Cell wall structural rigidity was also affected by drought conditions, substantially higher cellulose conversion rates were observed upon enzymatic saccharification of drought-treated samples with respect to controls. Both cell wall composition and the extent of cell wall plasticity under drought varied extensively among all genotypes, but only weak correlations were found with the level of drought tolerance, suggesting their independent genetic control.

### Genetic architecture of forage quality under well-watered and water deficit conditions

Among markers associated with forage quality under deferent irrigation episodes, a small number of the markers were in common between well-watered and water deficit conditions, while most of them responded independently to the treatments (Fig. 6), suggesting their independent genetic control. However, when phenotypic mean was used for GWAS, similar association patterns were found amongst energy-related traits, including DDM, TDN, ENE, ME, MEM, NEG and NEL, and traits of DRYMI, DMI1, RPV and RFQ (Fig. 7). The genetic responses to mean values of these traits may suggest common genetic bases among them. This is logical as these traits measure similar forage quality factors.

In the GWAS, we only find nine associated markers that have consistent effects across water deficit treatments (Fig. 8B). The rests were differentially associated with respective treatments. Interestingly, about 2 folds of markers were associated with mild water deficit compared to those identified by severe water deficit (Fig. 8B), suggesting that mild water deficit may trigger more genetic responses than the severe stress and the control. Drought tolerance is a complex trait and is affected by genetic and environmental interaction (GxE). This was dominated both at the trait level and for allelic effects of significantly associated causal variants. Therefore, we cannot directly address whether conditionally neutral alleles accumulate genetic variation at a faster rate than constitutively expressed genetic variation. For example, the number of significant markers were significantly reduced when severe water deficit applied compared to mild stress and well-watered control. This may indicate that the plants shut down some metabolic pathways to save energy for drought avoidance under severe drought stress.

### Putative candidate genes associated with forage quality

Among 23 annotated genes associated with forage quality traits, three genes were identified under well-watered condition (Table 3). The programmed cell death (PCD) protein was associated with DMI1, CP, RFQ, TDNL, NDFD48, IVTDMD-30 and −48. PCD in plants is a crucial component of development and defense mechanisms. PCD has multiple functions and regulates a complex molecular network in plant development (For review see Daneva et al., 2016). Its associations with multiple traits in the present study suggested that PCD involved in regulating forage quality in multiple ways. Reticulon-like protein B2 (RTNLB2) was associated with ash and TDNL. It has been reported that the RTNLB2 is located in endoplasmic reticulum and plays a role in regulating receptor transport to plasma membrane in Arabidopsis (Lee et al., 2011). Another putative candidate, cadmium/zinc transporting ATPase (cadA) was associated with NFC. The cadA is located in vacuole and involved in cadmium and zinc or cobalt transport and may have a detoxification function through a vacuolar sequestration in Arabidopsis (Mishra et al., 2017). Fourteen genes were identified under mild water deficit (Table 3). Of those, cytochrome P450 (CYP) was associated with ash, dNDF48, NDFD48, NCF and TDNL. P450 family protein is a large enzymatic protein family in plants and play a role in plant development and biotic and abiotic stresses responses (Xu et al., 2015 for review). UDP-glucosyltransferase (UGT) was associated with ash, NDFD48, NFC and TDNL. UGT plays a role in abscisic acid (ABA) homeostasis which regulates the plant response to environmental stresses such as drought cold and salinity (Dong et al., 2014). A RING zinc finger protein (RZFP) was associated with ash, NFC and TDNL under mild water deficit. It has been reported that overexpression of the Arabidopsis, AtRZFP enhanced salt and osmotic tolerance mediated by enhancing ROSs scavenging, maintaining Na^+^ and K^+^ homeostasis, adjusting the stomatal aperture to reduce water loss, and accumulating soluble sugars and proline to adjust the osmotic potential (Zang et al., 2016). An E3 ubiquitin protein ligase XBOS32 was associated with ash and NFC. There is evidence to suggest that E3 Ub-ligases can control protein turnover by modification of UPS-related proteins and contributes to nuclear proteome plasticity during plant responses to environmental stress signals (Serrano et al., 2018). An IQ calmodulin-binding motif protein was associated with ash. It has been reported that an IQ calmodulin-binding motif protein encoded by a gene of osa-mir369c classified as a small RNA family involved in impacting growth regulation under several environmental stresses such as temperature, drought and salinity in rice (Gao et al. 2010). The identification of calmodulin-binding proteins in the present study supports the assumption that this regulator is important player in response to abiotic stress through the calcium-signaling pathway (Ranty et al., 2006). Five genes were identified under severe water deficit (Table 3). Among them, the TIR-NBS-LRR protein was associated with ash and NFC. The plant TIR-NBS-LRR gene family contains a large class of disease resistance genes (DeYoung and Innes, 2006). The identification of the TIR-NBS-LRR genes in the present study suggests that it also involve in drought response in alfalfa. A similar finding of drought-related role for a NBS-LRR has also been reported in Arabidopsis where overexpression of the NBS–LRR gene ADR1 enhanced drought tolerance (Chini et al., 2004). It has been suggested that a signaling network exists between disease resistance and drought tolerance, and ADR1 may be involved in signal transduction in this network (Chini et al., 2004). An interferon-induced guanylate-binding protein (IIGBP) was associated with ash and NFC. The IIGBP is a GTPase induced by interferon and plays a role in directing inflammasome subtype-specific responses and their consequences for cell-autonomous immunity against a wide variety of microbial pathogens (Kim et al., 2016). A peroxidase family protein was associated with ash and NFC. The peroxidase responses are directly involved in the protection of plant cells against adverse environmental conditions. Several roles have been attributed to plant peroxidases in response to biotic and abiotic stresses. They may have a cell wall cross-linking activity during plant defense mechanisms (Chen et al., 2002). A RNA-binding (RRM/RBD/RNP motif) protein was associated with fat under severe water deficit. RNA-binding proteins (RBP) play important roles in post-transcriptional gene regulation. Recent investigation of plant RBPs demonstrated that, in addition to their role in diverse developmental processes, they are also involved in adaptation of plants to various environmental conditions (Lorkovic, 2009 for review). Although the remaining genes identified under water deficit do not have direct roles in stress responses, they involve in diverse processes in cell developments. For instance, Auxin-binding protein (ABP) was associated with ash, NFC and TDNL under mild water deficit. It has been suggested that ABP1 in Arabidopsis is involved in a broad range of cellular responses to auxin, acting either as the main regulator of the response, such as seen for entry into cell division or, as a fine-tuning device as for the regulation of expression of early auxin response genes (For review, see Tromas et al., 2010).

## Conclusion

In the present study, we evaluated 26 forage quality traits in a panel of 198 alfalfa accessions in the field trial under deficit irrigation gradience: well-watered, mild and severe water deficits. Phenotypic plasticities of forage quality traits were analyzed using the three irrigation treatments. Water deficit decreased fiber contents and enhanced energy-related traits. Higher correlations were found among the energy-related traits and the correlation coefficients increased when water deficit was applied. The highest correlation coefficient was obtained between RFQ and the quality mean, supporting that the RFQ is more accurate in estimating overall forage quality compared to the RFV.

Genetic architectures of phenotypic plasticities for 26 forage quality traits were investigated by GWAS under different irrigation treatments. Genetic markers associated with forage quality traits were identified and genetic regions responsive for the respective traits were compared. Similar regions were found between energy-related traits when mean values were used for GWAS. Significant markers associated with forage quality under water deficit were identified with highest significant marker-trait association in the mild stress. Only a small number of markers were commonly associated with all treatments. Most of the associated markers were independent to the different levels of water deficit treatments, suggesting genetic controls for forage quality traits were independent to the stress treatment. Although GWAS on forage quality have been reported, we are the first to address the genetic base of phenotypic plasticity of forage quality traits under water deficit. The information gained from the present study will be useful for the genetic improvement of alfalfa with enhanced drought/salt tolerance while maintaining forage quality.

## Acknowledgements

This work was supported partially by The United State Department of Agriculture NIFA_AFRP Grant Number 2015-70005-24071 and the Agriculture Research Service base fund. We thank Mrs. Martha Rivera for her technical assistance and data collection.

## Author Contributions

LXY planned and designed the research. BB and SF performed field work and forage quality tests. LXY, JH and SN wrote the manuscript.

## Abbreviations

ADF: Acid Detergent Fiber
DM: Dry Matter
DDM: Digestible Dry Matter
DMI: Dry Matter Intake using NDF
DMI1: Dry Matter Intake using NDF and NDFD
DDMI: Digestible Dry Matter Intake
ENE: Estimated Net Energy
IVDDM30: 30-hour In Vitro Digestible Dry Matter
IVDDM48: 48-hour In Vitro Digestible Dry Matter
ME: Metabolizable Energy
NEM: Net Energy for Maintenance
NDF: Neutral Detergent Fiber
dNDF: Digestible NDF
dNDF30: 30-hour Digestible NDF
dNDF48: 48-hour Digestible NDF
NDFD: NDF Digestibility
NDFD30: 30-hour NDFD
NEL: Net Energy for Lactation
NFC: Nonfibrous Carbohydrates
RFQ: Relative Forage Quality Index
RFV: Relative Feed Value Index
RUP: Rumen Undegradable Protein
TDN: Total Digestible Nutrients
TDNL: Total Digestible Nutrients for legume

